# A DNA tension-dependent tug-of-war between dimeric SMC motors governs loop extrusion directionality

**DOI:** 10.1101/2024.09.12.612694

**Authors:** Biswajit Pradhan, Adrian Pinto, Damla Tetiker, Takaharu Kanno, Gemma L. M. Fisher, Martin D. Baaske, Erin E. Cutts, Constantinos Chatzicharlampous, Herwig Schüler, Amar Deep, Kevin D. Corbett, Luis Aragon, Peter Virnau, Camilla Björkegren, Eugene Kim

**Author notes:** These authors contributed equally (Adrian Pinto, Damla Tetiker, Takaharu Kanno, Gemma L. M. Fisher).

## Abstract

Structural maintenance of chromosome (SMC) complexes organize and regulate genomes by extruding DNA loops. During loop extrusion, DNA can be reeled into the growing loop from one or both sides, generating distinct extrusion directionality states whose physical basis remains unclear. Here, we combine single-molecule analysis and molecular dynamics simulations to investigate loop extrusion directionality across SMC complexes. We show that dimeric Smc5/6 and Wadjet predominantly perform two-sided loop extrusion during initial loop growth, whereas monomeric condensin exhibits one-sided extrusion, consistent with a relationship between stoichiometry and extrusion directionality. Surprisingly, however, cohesin predominantly exhibits one-sided extrusion despite functioning as a dimeric complex. Notably, dimeric Smc5/6 and Wadjet progressively transition from two-sided to one-sided extrusion as loop growth matures. Simulations and force-dependent analysis reveal that loop extrusion directionality is governed by a tension-dependent transition in dimeric motors. Increasing DNA tension erodes two-sided extrusion near the motor stalling force, driving a tug-of-war regime in which competing motors transiently dominate one another. This transition promotes one-sided extrusion during mature Smc5/6 and Wadjet-mediated loop extrusion and places cohesin, which has a comparatively low stalling force (∼0.1 pN), constitutively near this regime. Together, our findings establish loop extrusion directionality as a dynamic emergent property governed by SMC stoichiometry, DNA tension, and stochastic motor competition.

## Introduction

Structural Maintenance of Chromosomes (SMC) complexes (henceforth referred to as SMCs), including condensin, cohesin and the Smc5/6 complex, play a critical role in organizing and maintaining genomes across all domains of life. At the molecular level, these complexes fold DNA into loops through a process known as loop extrusion. Loop extrusion has been directly visualized by *in vitro* single-molecule imaging for all eukaryotic SMC complexes (*1–7*) and for the prokaryotic SMC Wadjet complex (*8*). These experiments allow us to monitor the entire kinetics of loop extrusion, from the formation and enlargement of a loop, until its disruption, thus revealing important mechanistic details. One particularly intriguing and unresolved aspect of loop extrusion is the directionality by which SMCs reel DNA into a loop over time. Loop extrusion can be *two-sided* (*2, 3, 6, 8*), where SMCs reel in DNA simultaneously from both sides of a loop, or *one-sided* (*1*), where DNA is predominantly reeled in from only one side at a given time. One-sided extrusion can additionally exhibit direction switching, where the active side alternates over time; a behavior we refer to here as *one-sided with switching* (*9*). The directionality of loop extrusion is thought to have direct consequences for genome organization: two-sided extrusion is predicted to be more efficient at forming large domains and bringing distal genomic elements into proximity, while one-sided extrusion generates asymmetric loop structures with distinct implications for transcriptional regulation, sister chromatid cohesion, and DNA repair (*10*).

Despite its functional importance, the directionality of loop extrusion has remained difficult to resolve, with conflicting observations reported across species and even for the same complex under different conditions. For instance, yeast condensin has been observed to function as a strictly monomeric extruder and performs one-sided extrusion (*1*), whereas studies of human condensin have reported either exclusively one-sided extrusion (*11*) or a mixture of one-sided and two-sided extrusion behaviors (*4*). Similarly, while initial studies suggested two-sided extrusion by cohesin (*3*), recent work indicates one-sided extrusion with direction switching by single cohesin complexes (*9*). The directionality of loop extrusion by Smc5/6 has been a subject of debate, with conflicting reports of two-sided (*6*) and one-sided extrusion (*9*). Given that the physical principles governing loop extrusion directionality across SMC complexes remain poorly understood, these varying results are unsurprising.

In this study, we combine *in vitro* single-molecule fluorescence imaging and coarse-grained molecular dynamics simulations to systematically investigate the physical determinants of loop extrusion directionality across all known loop-extruding SMC complexes. We find that dimeric Smc5/6 and Wadjet predominantly undergo two-sided loop extrusion during early loop growth, whereas monomeric condensin exhibits one-sided extrusion, consistent with a relationship between stoichiometry and loop extrusion directionality. Surprisingly, cohesin also displays predominantly one-sided extrusion despite functioning as a dimeric complex. Notably, dimeric Smc5/6 and Wadjet progressively shift to one-sided extrusion during mature loop growth, indicating that stoichiometry alone cannot fully account for loop extrusion directionality. Using simulations and force-dependent analysis, we show that increasing DNA tension destabilizes coordinated activity between motors in dimeric SMC complexes, promoting competition between motors that results in one-sided extrusion and spontaneous direction switching. Such transition accounts for both the progressive shift toward one-sided extrusion by Smc5/6 and Wadjet during mature loop growth and cohesin’s predominantly one-sided extrusion behavior by placing cohesin constitutively near this mechanically sensitive regime. Together, our results provide a unified physical framework for understanding loop extrusion directionality across SMC complexes, in which directional states emerge from the interplay between SMC stoichiometry, DNA tension, and stochastic motor competition.

## Results

### Assessing loop extrusion directionality across SMC complexes

To investigate the directionality of DNA loop extrusion by SMC complexes, we employed *in vitro* single-molecule fluorescence imaging to monitor loop extrusion events in real time. In this assay, both ends of a DNA molecule are anchored to a passivated glass surface and stained with the intercalating dye SYTOX Orange (Fig. 1a). DNA molecules are stretched via buffer flow during deposition to approximately 30% of their contour length. When an SMC complex binds and extrudes a loop, the loop appears as a bright spot along the DNA and gradually expands, as depicted in the example kymograph in Fig. 1b. The growing loop is flanked by two shrinking DNA segments of lengths I and II on either side. A typical loop extrusion event proceeds through two phases: an initial growth phase during which the loop rapidly expands and reaches a plateau, followed by a mature phase characterized by slower dynamics with alternating loop shrinkage and regrowth (Fig. 1c).

**Figure 1.**
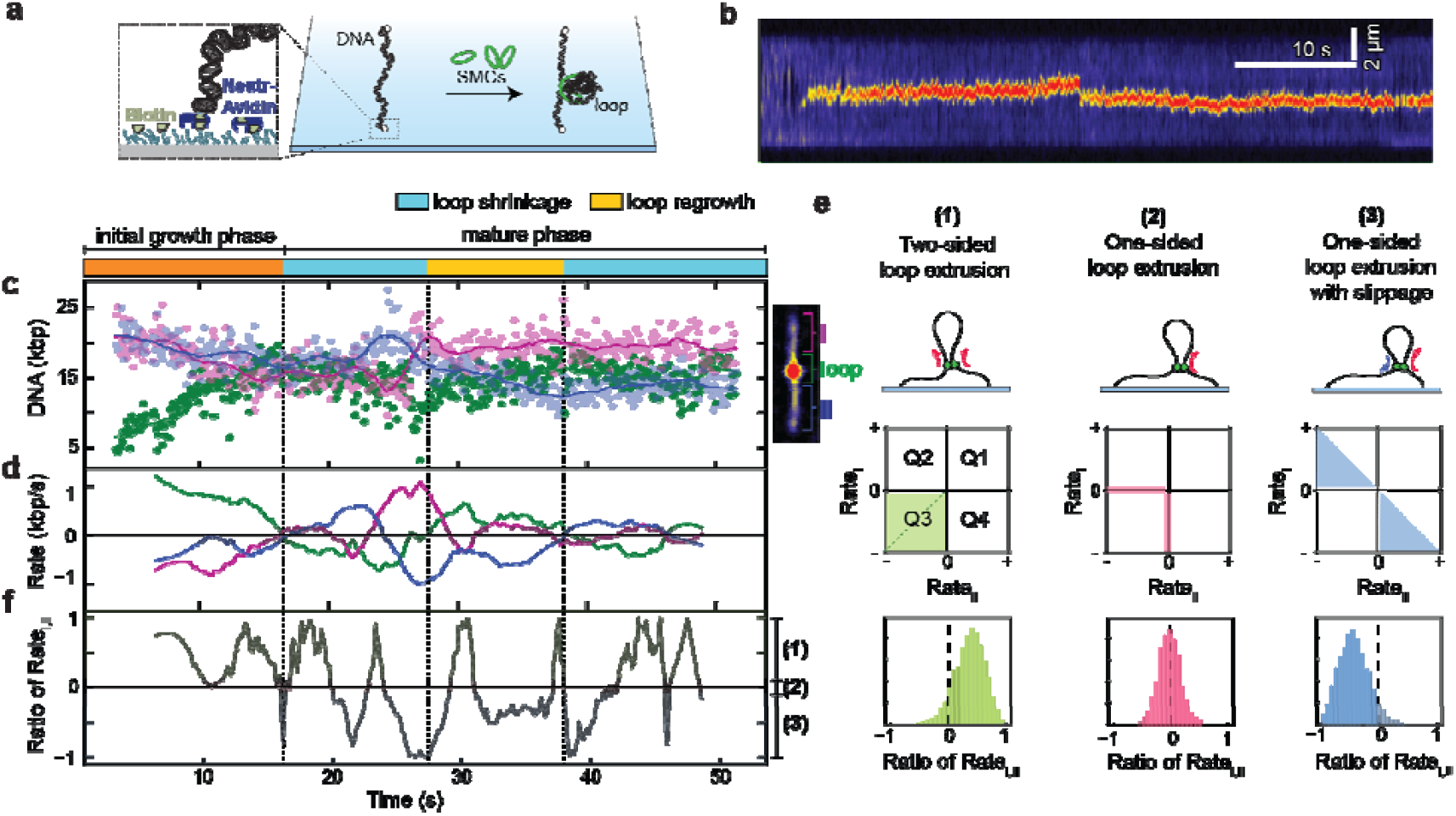
Evaluating loop extrusion directionality by SMC complexes. (a) Schematic of a flow cell containing surface-anchored λ-DNA used for monitoring DNA loop extrusion by SMCs. (b) Representative kymograph of a DNA molecule with a loop extruded by Smc5/6. (c) DNA lengths calculated from the kymograph in b for the two DNA segments flanking the loop (I, II) and the loop itself, as indicated in b. (d) The rates of DNA length change for the traces shown in c, extracted via linear fitting over 5-second intervals using a running time window. (f) The ratios of the rates, Rate_small_/Rate_large_, where Rate_small_ (Rate_large_) correspond to the smaller (larger) absolute values of Rate_I_ and Rate_II_, respectively. (e) Illustration showing three loop extrusion directionality classes classes(top), their corresponding quadrant plots of Rate_I_ versus Rate_II_ (middle), and the resulting ratio of the rates (bottom). The green diagonal in the quadrant plot denotes strictly symmetric two-sided extrusion (Rate_I_ = Rate_II_).

To quantify loop extrusion directionality, we calculated the rates of change of DNA segments I and II (Rate_I_, Rate_II_) throughout extrusion (Fig. 1d; Methods). Importantly, our analysis evaluates extrusion directionality continuously over time, allowing tracking of individual complexes when dynamically transitioning between distinct extrusion states during a single loop extrusion event. This time-resolved approach therefore captures both stable and transient directional behaviors and weights directionality distributions according to the time spent in each state. The instantaneous extrusion states are most directly visualized using a quadrant plot of Rate_I_ versus Rate_II_ (Fig. 1e; Fig. S1). Time points in which both rates are simultaneously negative (Q3) correspond to two-sided extrusion, whereas points along a single negative axis indicate strictly one-sided extrusion. Points with one negative and one positive rate reflect one-sided extrusion with DNA slippage from the passive side. To enable quantitative comparison across complexes and maintain consistency with prior literature, we additionally summarize these distributions using the ratio-of-rates metric (Rate_small_/Rate_large_; Fig. 1f; Fig. S1). Positive ratios (> 0.1) indicate two-sided extrusion, ratios near zero (between −0.1 and +0.1) indicate strictly one-sided extrusion, and negative ratios (< −0.1) indicate one-sided extrusion with slippage.

### Initial loop extrusion symmetry differs across SMC complexes but converges to one-sided growth during loop maturation

We first systematically characterized the directionality of loop extrusion for all four SMC complexes using the time-resolved framework described above (Fig. 2). For Smc5/6 (N = 72), the quadrant plot during the initial growth phase showed a broad distribution of points centered in Q3, indicating that both Rate_I_ and Rate_II_ are simultaneously negative; that is, both flanking DNA segments shorten together, consistent with two-sided extrusion (Fig. 2a,b). This two-sided behavior was most pronounced during initial growth (52% two-sided fraction; Fig. 2g), whereas during the mature phase the fraction of two-sided events decreases and one-sided extrusion with slippage became predominant (Fig. 2c–g, Fig. S1). A time-dependent analysis further revealed a gradual decline of the two-sided fraction from ∼65% to ∼34% over the course of initial growth (Fig. S2a,b). Importantly, two-sided extrusion remained prominent also at 10 ms temporal resolution (63% two-sided during initial growth; Fig. S2c–j), ruling out the possibility that the apparent two-sided signal reflects unresolved asynchronous one-sided extrusion at faster timescales. The two-sided fraction was also robust across different time-segment lengths used for rate calculation (Fig. S2k). Additional control analyses confirmed that the 5-second window used throughout this study provides reliable segment-rate measurements without substantial contributions from short-timescale DNA fluctuations (Fig. S2k-m).

**Figure 2.**
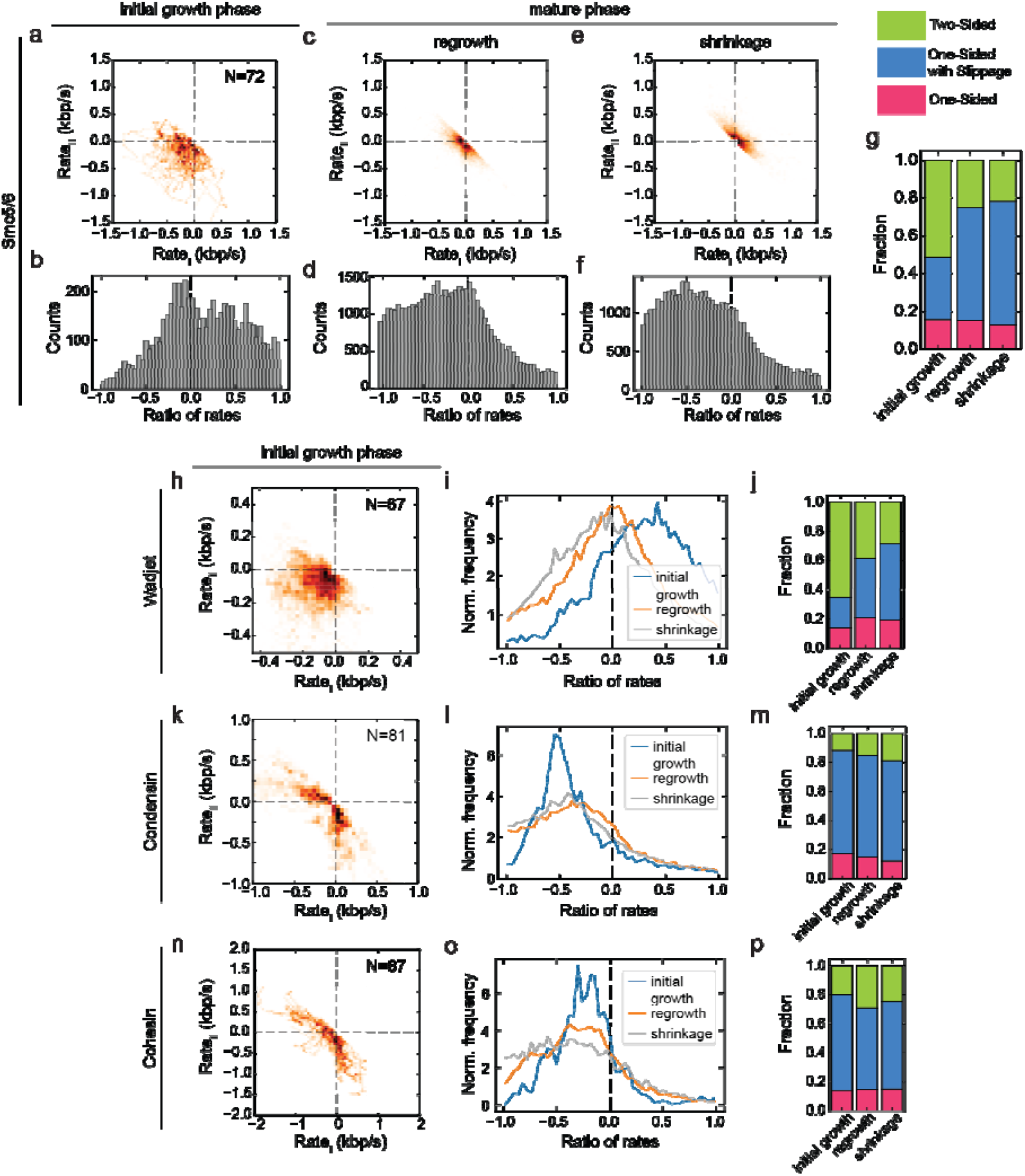
Loop extrusion directionality across SMC complexes. (a,c,e) Quadrant plots showing the rates of DNA length change in the two DNA segments flanking the loop (Rate_I_, Rate_II_), and the corresponding ratios of rates (b,d,f ), for Smc5/6 during the phases of initial loop growth (a,b), mature loop growth (c,d), and loop shrinkage (e,f). (g) Fractions of extrusion symmetry classes observed during the three phases. (h,k,n) Quadrant plots of Rate_I_ versus Rate_II_, with the corresponding rate-ratio distributions (i,l,o) and fractions of extrusion directionality classes (j,m,p) for Wadjet (h-j), yeast condensin (k-m) and human cohesin (n-p).

Wadjet (N = 67) showed a similar pattern, with strong two-sided extrusion during initial growth (65%; Fig. 2h–j), followed by predominantly one-sided extrusion with slippage during the mature phase (Fig. 2i,j, Fig. S3a-d). In contrast, condensin (N = 81) and cohesin (N = 87) showed negligible Q3 occupancy and ratio-of-rates distributions strongly skewed below zero throughout all phases (Fig. 2k–p, Fig. S3e-l), indicating that both complexes predominantly exhibit one-sided extrusion throughout loop growth. Taken together, Smc5/6 and Wadjet predominantly exhibit two-sided extrusion during early loop growth, whereas one-sided extrusion becomes a shared feature of all SMC complexes in the mature phase.

### Stoichiometry correlates with, but does not fully determine, loop extrusion directionality

The distinct extrusion symmetries observed across SMC complexes prompted us to ask whether loop extrusion directionality is determined by the stoichiometry of the actively extruding complexes. To address this, we fluorescently labeled each complex’s kleisin subunit, which is present at a stoichiometry of one kleisin per SMC motor subunit homo/heterodimer – and performed photobleaching step analysis during active loop extrusion events (Fig. 3a–e). Because fluorescent tagging could potentially perturb loop extrusion dynamics, we first assessed its effect on Smc5/6 behavior. Tagged Smc5/6 complexes exhibited increased loop mobility and a higher tendency for DNA slippage compared to untagged wild-type complexes, shifting extrusion behavior toward more one-sided, slippage-dominated states (Fig. S4).

**Figure 3.**
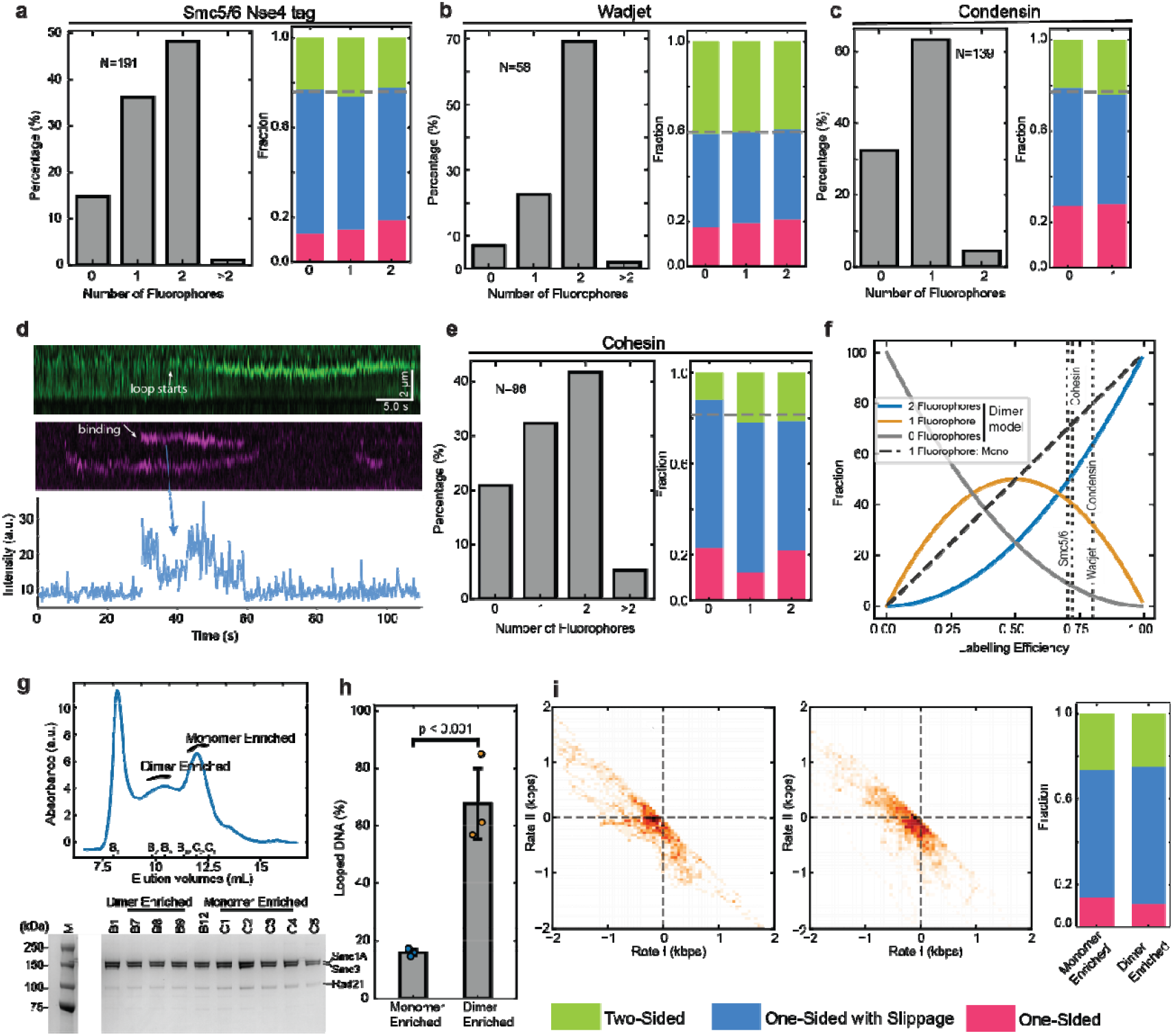
Stoichiometry of SMC complexes during loop extrusion and its relationship to extrusion directionality. (a–c) Fluorophore stoichiometry and extrusion directionality of Smc5/6 (a), Wadjet (b), and condensin (c). Histograms (left) show the distribution of fluorophore counts per loop extrusion event, and stacked bar plots (right) show the fractions of two-sided, one-sided with slippage, and strictly one-sided extrusion for subpopulations grouped by fluorophore count. (d) Representative loop extrusion event by fluorescently labeled cohesin showing the DNA channel (top, green), cohesin-Alexa647 channel (middle, magenta), and fluorescence intensity trace showing two step bleaching event (bottom, blue). The transient increase in intenisity following the first bleaching reflects binding of additional non-extruding cohesin molecules to the DNA, rather than recruitment to the extruding complex. (e) Fluorophore stoichiometry and extrusion directionality for cohesin labeled with Alexa647. (f) Theoretical fluorophore distributions for dimeric and monomeric models as a function of labeling efficiency. Vertical dashed lines indicate the measured labeling efficiencies for each SMC complex. (g) Size exclusion chromatography profile of unlabeled cohesin, resolved on a Superose 6 column, showing distinct dimer-enriched and monomer-enriched peaks (top), and SDS-PAGE analysis of fractions collected from each peak (bottom). (h) Loop extrusion efficiency for monomer-enriched (Peak 2) and dimer-enriched (Peak 1) cohesin fractions at an equivalent concentration of 4 nM monomeric units. (i) Quadrant plots of Rate I versus Rate II for loop extrusion events from the monomer-enriched (left) and dimer-enriched (middle) cohesin fractions, and the corresponding directionality fractions (right).

Photobleaching analysis of Smc5/6 labeled at Nse4 via SNAP-tag with Alexa647 (DoL = 70%; N = 191) revealed that the fluorophore count distribution was consistent with a dimeric model, with the majority of events showing one or two fluorophores (Fig. 3a, Fig. S5a). Wadjet labeled at JetA with JF650 (DoL = 80%; N = 58) showed a similar distribution, again supporting dimeric extrusion (Fig. 3b). In contrast, condensin labeled at Ycs4 via ybbR-tag with ATTO647N (DoL = 80%; N = 139) predominantly showed single-step bleaching, consistent with extrusion by a monomeric complex (Fig. 3c, Fig. S5b). Cohesin labeled at Rad21 via a C-terminal ybbR tag conjugated to Alexa647 (DoL=72%) similarly displayed predominantly one- and two-step bleaching during active loop extrusion events, indicating that the actively extruding unit contains two cohesin complexes (Fig. 3d–f, Fig. S5c-d). Consistently, fluorescence intensity analysis of surface-adsorbed cohesin molecules revealed a distinct two-fluorophore population (∼20%), independently supporting the presence of dimeric cohesin under loop extrusion conditions (Fig. S5h,i). Together, these fluorophore distributions are consistent with dimeric extrusion by Smc5/6, Wadjet, and cohesin, and monomeric extrusion by condensin (Fig. 3f).

Because labeling efficiency was below 100%, fluorophore count distributions alone cannot unambiguously distinguish a purely dimeric population from a mixture of monomeric and dimeric extruders. If the observed extrusion asymmetries reflected differences in extruder stoichiometry, then complexes with different fluorophore occupancies (0, 1, or 2 fluorophores) should exhibit different extrusion behaviors. However, the fractions of two-sided, strictly one-sided, and one-sided-with-slippage extrusion remained unchanged across all fluorophore occupancy classes (Fig. 3a–e, Fig. S6). Thus, while tagging can modestly alter the overall extrusion behavior (Fig. S4), extrusion directionality is largely independent of fluorophore occupancy, further supporting the conclusion that the observed asymmetric extrusion behaviors arise from complexes with the same underlying stoichiometry.

To further probe the relationship between cohesin stoichiometry and extrusion directionality, we separated monomer-enriched and dimer-enriched fractions by size exclusion chromatography (SEC; Fig. 3g). SDS-PAGE confirmed the protein composition of each eluted fraction. Loop extrusion assays performed at equivalent monomeric concentrations (4 nM) revealed substantially higher loop extrusion activity in the dimer-enriched fraction, with ∼60% of DNA molecules forming loops compared to ∼15% for the monomer-enriched fraction (p < 0.001; Fig. 3h), supporting the idea that dimeric cohesins represent the active loop-extruding form. Notably, directionality analysis of loop extrusion events from both monomer-enriched and dimer-enriched fractions showed similar one-sided extrusion behavior (Fig. 3i).

Together, these results establish a broad correlation between stoichiometry and initial extrusion symmetry: Smc5/6 and Wadjet, which predominantly exhibit two-sided extrusion during early loop growth, extrude DNA as dimers, whereas condensin, which exhibits strictly one-sided extrusion, functions primarily as a monomeric extruder. However, cohesin presents a notable exception to this trend. Despite functioning as a dimeric complex, cohesin predominantly exhibits one-sided extrusion behavior, suggesting that dimerization alone is insufficient to maintain stable two-sided extrusion.

### Asymmetric loop extrusion emerges from symmetric dimeric motors

The observation that Smc5/6 and Wadjet progressively transition toward one-sided extrusion, whereas cohesin predominantly exhibits one-sided extrusion despite all three functioning as dimeric complexes, raised a central question: how can symmetric dimeric motors generate one-sided extrusion?

To address this question, we performed coarse-grained molecular dynamics simulations of dimeric loop extrusion using a minimal two-ring “handcuff” model for SMC complexes coupled to a bead-spring representation of DNA (Fig. 4a) (10,11). In this model, the two rings act as independent DNA-reeling motors exerting equal extrusion forces, representing an idealized symmetric dimeric extruder. Simulations produced progressive loop growth followed by stalling at larger loop sizes, closely resembling the experimentally observed transition from initial growth to mature extrusion behavior (Fig. 4b,c). During loop growth, the DNA segments flanking the loop (I and II) shortened simultaneously, consistent with two-sided extrusion by Smc5/6 and Wadjet.

**Figure 4.**
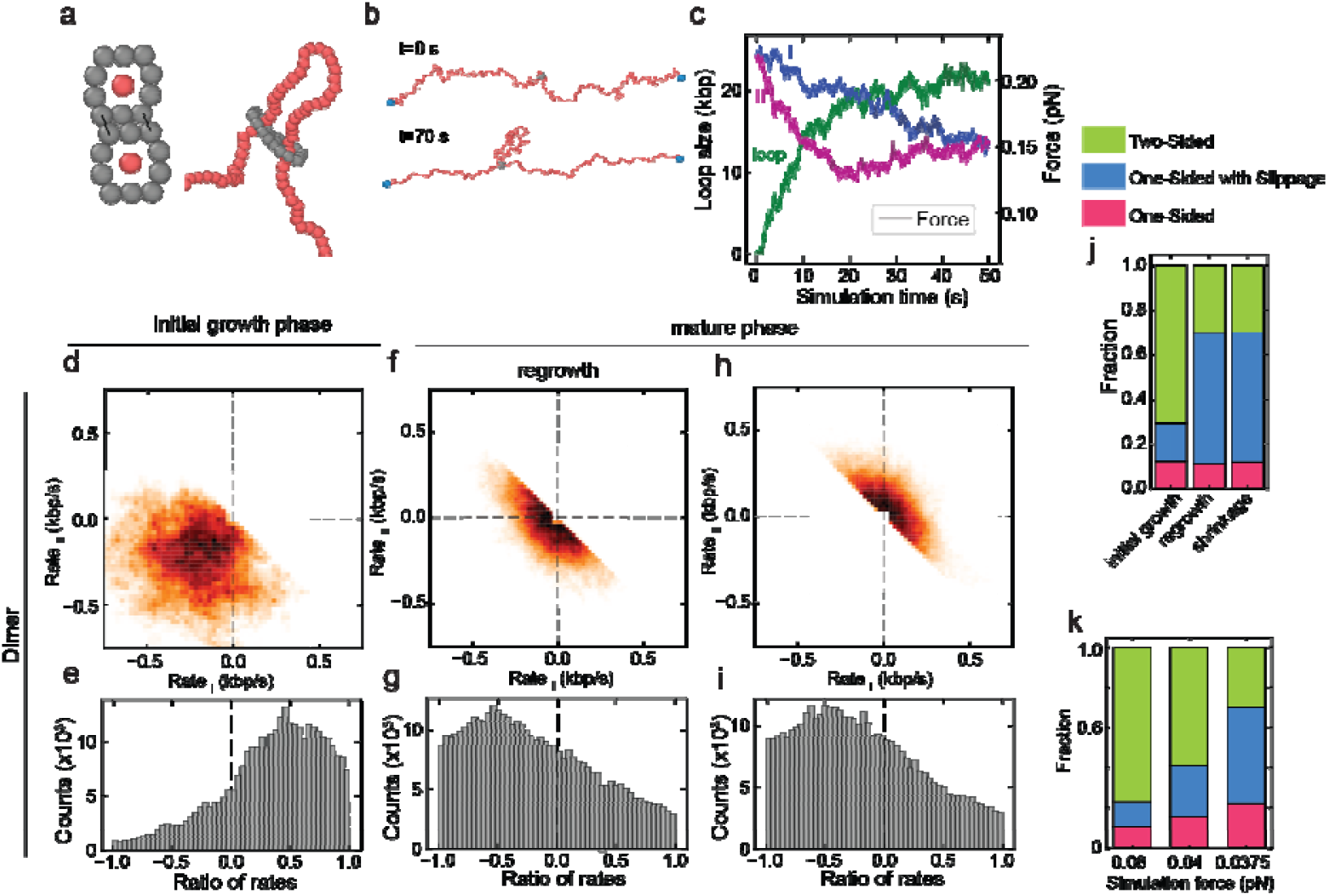
One-sided loop extrusion emerges inherently from symmetric dimeric motors. (a) Schemati of the coarse-grained simulation model showing the two-ring “handcuff” structure (grey) representing a dimeric SMC complex, and the bead-spring polymer (red) representing DNA. (b) Snapshots of a simulated loop extrusion event at t = 0 s (top) and t = 70 s (bottom), illustrating progressive loop formation by the dimeric motor. (c) Representative time traces of DNA segment sizes (I, blue; II, green; loop, magenta) and DNA tension outside the loop (grey) over the course of a simulation. (d–j) Directionality analysis of simulated dimeric motor loop extrusion (N = 22). Quadrant plots of Rate_I_ versus Rate_II_ (d,f,h), corresponding rate-ratio distributions (e,g,i), and the resulting fractions of extrusion symmetry classes (j) during the initial growth phase (d,e), mature growth phase (f,g), and mature shrinkage phase (h,i). (k) Extrusion directionality fractions for simulations performed at three different extrusion forces (0.06, 0.04, and 0.0375 pN). The higher force regime (0.06pN) generate stalling forces comparable to those measured experimentally for Smc5/6 and Wadjet, whereas the lowest force regime (0.0375pN) is comparable to the stalling forces of cohesion.

Surprisingly, despite the complete symmetry of the model, simulated dimeric motors frequently exhibited transient one-sided extrusion. Although both motors exerted identical forces calibrated to experimentally measured SMC stalling force regimes (Fig. S7a–c; Methods), the DNA reeling rates of the two motors fluctuated independently over time (Fig. 4c-e). Consistent with this prediction, single-molecule translocation analysis of individual Smc5/6 complexes revealed substantial variability in DNA reeling rates both between molecules and within individual trajectories (Fig. S8). As a result, extrusion trajectories often appeared one-sided even though both motors remained active. The resulting simulated kymographs, DNA segment trajectories, and rate distributions closely resembled the experimentally observed extrusion patterns of Smc5/6 and Wadjet (Fig. S7; Fig. 4c–e). Notably, simulations further reproduced the experimentally observed progressive transition toward one-sided extrusion during mature loop growth (Fig. 4f-j). Even in the absence of imposed asymmetry or explicit switching rules, coordinated two-sided extrusion became increasingly unstable as extrusion progressed, leading to progressively more asymmetric extrusion behavior.

We reasoned that this progressive destabilization may arise from increasing DNA tension during loop growth. In the simulations, loop extrusion begins under low-tension conditions and progressively generates increasing opposing DNA tension as the loop enlarges, analogous to the experimental single-molecule assay. To test whether the balance between motor force and opposing DNA tension influences the stability of coordinated two-sided extrusion, we performed simulations across a range of extrusion forces chosen relative to the experimentally measured stalling-force regimes of Smc5/6, Wadjet and cohesin (Methods; (*12*)). Lowering the extrusion force progressively reduced the fraction of two-sided extrusion events during the initial growth, leading to increasingly one-sided extrusion (Fig. 4k). Notably, forces approaching the cohesin stalling-force regime produced strongly asymmetric extrusion dynamics despite the fully symmetric motor architecture. These findings suggest that lower-force motors operate near a mechanically unstable regime in which coordinated two-sided extrusion becomes difficult to maintain.

To model monomeric extrusion, we eliminated force generation from one of the two rings in the handcuff model (Fig. S7g,h,k,n). This primarily resulted in DNA translocation without stable loop extrusion, consistent with previous observations for individual Smc5/6 complexes(*6, 9*) (Fig. S8). Introducing a weak anchoring interaction restored stable loop extrusion and produced predominantly one-sided extrusion (Fig. S7i-q).

Together, these simulations demonstrate that one-sided loop extrusion can emerge from both monomeric and dimeric motor architectures through distinct physical mechanisms. Whereas monomeric extruders generate inherently one-sided extrusion, symmetric dimeric motors can also develop persistent asymmetry despite retaining two active motors.

### DNA tension drives a stochastic tug-of-war regime in dimeric SMC complexes

Our simulations suggested that increasing DNA tension destabilizes coordinated two-sided extrusion in symmetric dimeric motors. We therefore next tested experimentally whether loop extrusion directionality is directly regulated by DNA tension. To test this idea, we quantified loop extrusion directionality as a function of DNA tension using the known force-extension relation of DNA (15) (Fig. 5a–h). For both Smc5/6 and Wadjet, extrusion directionality strongly depended on DNA tension. At low tensions, both complexes predominantly displayed two-sided extrusion with ratio-of-rates values near unity (Fig. 5a–b). As DNA tension increased, the ratio-of-rates sharply decreased, indicating a progressive transition toward one-sided extrusion. This transition occurred near, but slightly below, the motor stalling regime (stalling tensions of ∼0.1 pN vs 0.25 pN for Smc5/6 and Wadjet) and coincided with a pronounced reduction in loop growth rate. In contrast, condensin exhibited predominantly one-sided extrusion across the full tension range with little dependence on DNA tension (Fig. 5c), consistent with its monomeric extrusion architecture.

**Figure 5.**
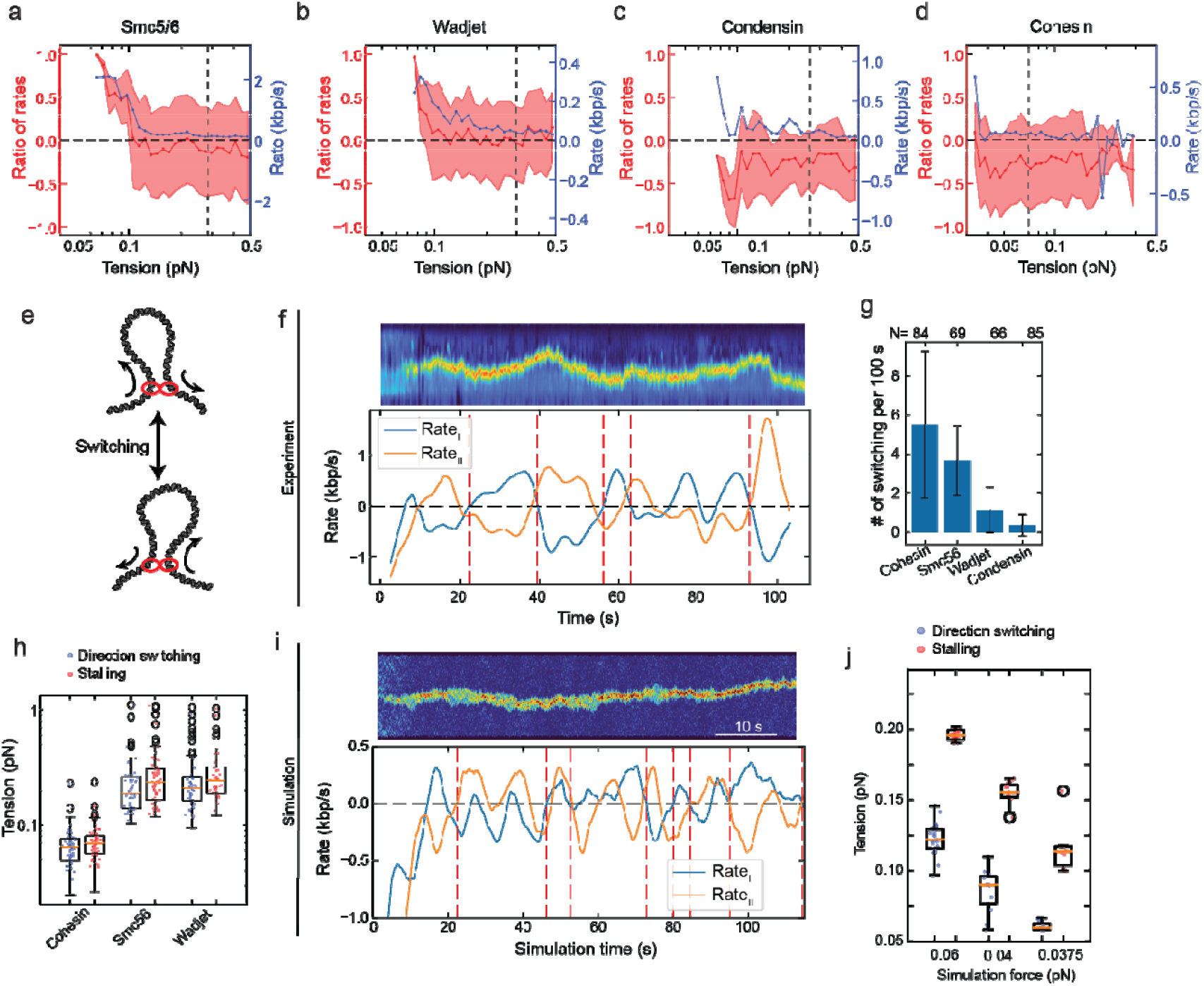
DNA tension drives a stochastic tug-of-war regime in dimeric SMC complexes. (a–d) Ratio-of-rates (red, left axis, mean ± s.d.) and loop extrusion rate (blue, right axis) plotted as a function of DNA tension outside the loop for Smc5/6 (a), Wadjet (b), condensin (c), and cohesin (d). Dashed vertical lines in (a-d) indicate the average stalling force for the corresponding SMC complex. (e) Schematic illustrating direction switching during one-sided loop extrusion by a dimeric extruder, accompanied by slippage from the non-extruding side. (f) Representative experimental kymograph (top) and corresponding time traces of Rate_I_ (blue) and Rate_II_ (orange) showing repeated direction-switching events (red dashed lines) during loop extrusion by a dimeric Smc5/6 complex. (g) Direction-switching frequency, quantified as the number of switching events per 100 s for cohesin (N = 84), Smc5/6 (N = 69), Wadjet (N = 66), and condensin (N = 85). Error bars indicate standard deviation. (h) Box plots of DNA tension at the first direction-switching event (blue) and at stalling (red) for cohesin, Smc5/6, and Wadjet. (i) Simulated kymograph (top) and Rate_I_-Rate_II_ time traces (bottom) for a symmetric dimeric motor, showing spontaneous direction-switching events (red dashed lines) emerging without any explicit switching mechanism. (j) Box plots of switching tension from simulations performed at three different extrusion forces of 0.06, 0.04, and 0.0375 pN). These conditions generate stalling-forces comparable to the experimentally measure stalling-force regimes for Wadget, Smc5/6, and cohesin, with higher-force regimes (0.06 pN, 0.04 pN) more closely reflecting Wadjet and Smc5/6, whereas the lowest-force (0.0375 pN) regime more closely reflecting cohesin.

Cohesin similarly displayed predominantly one-sided extrusion even at low experimentally accessible tensions (Fig. 5d), suggesting that cohesin operates near the regime where stable two-sided extrusion is difficult to maintain. Consistently, simulations showed that reducing the extrusion force of dimeric motors from the Smc5/6- and Wadjet-like regime (0.06 pN) toward cohesin-like regime (0.0375 pN) progressively decreased the fraction of two-sided extrusion states, shifting the system toward predominantly one-sided extrusion (Fig. 4k). Notably, externally applied buffer flow promoted two-sided extrusion by cohesin (Fig. S10a-h), indicating that relatively small external forces can rebalance motor activity.

If both motors remain active near stalling, transient dominance of one motor over the other should generate spontaneous switching between extrusion directions. To test this prediction, we quantified direction switching during loop extrusion by monitoring sign changes in Rate_I_ and Rate_II_ throughout individual extrusion events (Fig. 5e,f; Fig. S9i-m; Methods). Smc5/6 exhibited frequent switching between extrusion directions, with switches occurring on average every ∼27 s, whereas Wadjet switched substantially less frequently (∼80 s between switches; Fig. 5g). In contrast, condensin rarely switched direction (∼278 s between switches), consistent with predominantly one-sided extrusion by a monomeric motor. Cohesin exhibited the most frequent switching behavior (∼18 s between switches), consistent with frequent loss of directional balance between the two motors near stalling.

Importantly, the first switching events for Smc5/6 and Wadjet occurred at DNA tensions near, but lower than, the respective motor stalling forces (Fig. 5h; Fig. S9k). Cohesin, by contrast, exhibited switching under essentially all experimentally accessible conditions, consistent with its comparatively low stalling tension (∼0.07 pN versus ∼0.2 pN for Smc5/6 and Wadjet; Fig. 5h). Consistent with this interpretation, coarse-grained simulations reproduced both the experimentally observed switching behavior and its force dependence without incorporating any dedicated switching mechanism (Fig. 5i,j, Fig. S9l,m). Instead, switching emerged naturally from competition between two motors operating near their stalling forces.

Together, these results demonstrate that increasing DNA tension drives dimeric SMC complexes into a stochastic tug-of-war regime characterized by one-sided extrusion, DNA slippage, and spontaneous direction switching. Cohesin’s predominantly one-sided extrusion behavior can therefore be explained by its comparatively low stalling force, which places cohesin constitutively near this regime even under low experimental tensions.

## Discussion

### A unified framework: stoichiometry and tension govern loop extrusion directionality

Our results support a unified framework in which loop extrusion directionality reflects the mechanical state of interacting SMC motors. Dimeric complexes transition from coordinated two-sided extrusion at low tension to a tug-of-war regime characterized by one-sided extrusion, slippage, and direction switching near stalling, whereas monomeric condensin remains predominantly one-sided independent of tension. Our molecular dynamics simulations show that this full spectrum of behaviors emerges naturally from basic motor physics. Simulations using two identical motors reproduced coordinated two-sided extrusion at low tension, one-sided extrusion with slippage and spontaneous direction switching near the stalling tension. The transition from two-sided to one-sided extrusion therefore emerges naturally from competition between mechanically coupled motors without requiring an explicit switching mechanism.

### Cohesin extrudes DNA loop as a dimer

One of the major findings of this study is that cohesin extrudes DNA loops as a dimer despite predominantly exhibiting one-sided extrusion behavior. This conclusion is supported by two independent observations: photobleaching stoichiometry during active extrusion, which is consistent with a dimeric but not monomeric model (Fig. 3d–f), and the substantially higher loop extrusion activity of the dimer-enriched cohesin fraction compared to the monomer-enriched fraction after SEC separation (Fig. 3g,h). Furthermore, both fractions exhibited indistinguishable extrusion directionality profiles (Fig. 3i), indicating that the residual activity in the monomer-enriched fraction likely arises from residual or newly formed dimers rather than genuinely monomeric extrusion.

Our findings are consistent with previous reports of two-step photobleaching during cohesin-mediated loop extrusion (Kim et al.(*3*)) and with recent in vivo imaging studies identifying a major dimeric population of chromatin-bound cohesin in post-replicative cells (Ochs et al.(*13*)). Together, these observations support a model in which dimeric cohesin serves as the primary functional unit both *in vitro* and *in vivo*.

This interpretation differs from earlier reports describing one-sided extrusion by single cohesin complexes (*2, 9*). However, cohesin stoichiometry measurements during loop extrusion are technically challenging and sensitive to experimental conditions such as laser power, including fluorophore photobleaching prior to extrusion onset and the relatively long delay between DNA binding and loop initiation observed for cohesin (Fig. S5e-g). Such factors may contribute to differences in the reported stoichiometry of cohesin-mediated loop extrusion across studies.

### Reconciling discrepancies in reported extrusion directionality

Our results also help reconcile discrepancies between previous studies of SMC directionality. In particular, our observations differ from Barth et al. (*9*), who reported predominantly asymmetric extrusion for all eukaryotic SMC complexes. One likely explanation is the difference in analytical approaches. Our analysis quantifies extrusion directionality continuously throughout individual loop extrusion events, thereby resolving dynamic transitions between two-sided and one-sided behaviors. By contrast, phase-based classifications may underrepresent transient two-sided extrusion during early loop growth, particularly because all dimeric complexes progressively lose coordinated two-sided extrusion near stalling.

Because loop extrusion directionality reflects the mechanical state of the extruding complex, it is also expected to respond sensitively to perturbations that alter DNA tension or motor coordination. Consistent with this idea, we find that extrusion directionality is influenced by multiple physicochemical parameters, including protein tagging, ionic conditions, DNA staining, and externally applied forces (see Supplementary Notes 1-4). Notably, externally applied perpendicular side-flow promoted two-sided extrusion by cohesin (Fig. S10a) and similarly stabilized in simulations two-sided extrusion by counteracting DNA slippage (Fig. S10b-h), supporting the idea that relatively small external forces can shift extrusion directionality in mechanically sensitive regimes. Together, these observations suggest that loop extrusion directionality is highly sensitive to both mechanical and experimental perturbations.

### Biological implications of tension-dependent extrusion directionality

Our results suggest that loop extrusion directionality is not a fixed property of SMC complexes but instead depends on the physical environment of the DNA substrate. In vivo, local DNA tension is shaped by transcription, replication, chromatin compaction, DNA supercoiling, and other topological constraints. Because these processes act locally and dynamically, the mechanical loads experienced by an extruding complex are expected to vary substantially across chromosomal contexts.

Under such conditions, the ability to transition between coordinated two-sided extrusion and one-sided extrusion with direction switching may provide functionality beyond that achievable by either mode alone. Coordinated two-sided extrusion efficiently compacts DNA by simultaneously reeling in both flanks of a loop. As tension builds, however, further compaction becomes progressively less favorable. Rather than terminating extrusion at this point, a transition to a competitive regime can allow continued loop translocation through one-sided extrusion and direction switching. Coordinated and competitive extrusion may therefore represent complementary operational states of the same molecular machine: one optimized for efficient DNA compaction and the other for repositioning loops along DNA once further compaction becomes limited. In this framework, DNA tension acts as a physical regulator that shifts SMC complexes between these regimes.

Smc5/6 provides a particularly clear example of this behavior. Under low tension, Smc5/6 predominantly exhibits coordinated two-sided extrusion, whereas increasing DNA tension shifts the complex toward asymmetric extrusion and frequent direction switching. Given the association of Smc5/6 with supercoiled DNA, replication-associated structures, and sites of DNA damage, it is tempting to speculate that similar transitions may occur in vivo as local mechanical conditions change. Such transitions could allow Smc5/6 not only to compact DNA efficiently under favorable conditions but also to continue loop translocation once the limits of coordinated compaction have been reached.

Wadjet may similarly exploit coordinated two-sided extrusion under low mechanical load while transitioning toward competitive extrusion as tension increases. Such flexibility could support efficient DNA capture and compaction while preserving the ability to reposition loops across a broader range of substrate geometries and mechanical environments. Whether these transitions contribute to substrate discrimination between plasmid and chromosomal DNA remains an intriguing possibility for future investigation.

Cohesin appears to operate closer to the transition between coordinated and competitive regimes, consistent with its relatively low stalling force. Because cohesin-mediated loop extrusion often functions to position loops relative to genomic landmarks rather than simply maximize compaction, the ability to continue loop translocation after the limits of coordinated compaction have been reached may facilitate loop repositioning and boundary searching during genome organization. Recent work has further shown that NIPBL exchange can influence cohesin direction switching, suggesting that regulatory mechanisms may tune switching dynamics within this mechanically sensitive regime (*9*).

By contrast, condensin is monomeric and constitutively one-sided, placing it outside the dimeric tug-of-war regime. Its predominantly one-sided extrusion behavior is consistent with models in which chromosome compaction emerges from highly processive asymmetric extrusion coupled with collective condensin activity. In this context, condensin traversal and the formation of Z-loops may provide an alternative mechanism for maintaining chromosome-scale compaction without requiring transitions to coordinated two-sided extrusion (*14*).

Collectively, these observations support a view in which coordinated and competitive extrusion represent complementary operational states of the same molecular machine. Different SMC complexes may favor different regions of the extrusion landscape, with DNA tension shifting their behavior between coordinated and competitive regimes. In this framework, direction switching is not simply a deviation from two-sided extrusion but a physically regulated behavior that enables continued loop translocation even after the limits of efficient DNA compaction have been reached.

### Conclusion

Together, our findings establish loop extrusion directionality as a dynamic emergent property governed by SMC stoichiometry, DNA tension, and stochastic motor competition. By combining comparative single-molecule analysis with molecular dynamics simulations across all known loop-extruding SMC complexes, we show that one-sided extrusion and direction switching can arise naturally from mechanically unstable coordination between dimeric motors near stalling. These results reveal how distinct loop extrusion behaviors can emerge from common underlying physical principles operating under different mechanical regimes. More broadly, our results suggest that local chromosome environments, through their effects on DNA tension, may actively tune loop extrusion dynamics to regulate genome organization *in vivo*.

## Acknowledgments

We thank Gabriel Maul for the production of biotinylated λ-DNA. AP and PV acknowledge fruitful discussions on simulations with Apratim Chatterji.

## Funding

This work was supported by Max Planck Society (EK), European Research Council Starting Grant 101076914 (EK), Hessian Ministry of Science and Art MSCA Grant (BP), Swedish Cancer foundation, Swedish research council, and Centre for Innovative Medicine (CIMED) (CB), National Institutes of Health R35 GM144121 (KDC), Medical Research Council UKRI MC-A652-5PY00 (L.A., and E.E.C). EK, DT, AP and PV are grateful to the Deutsche Forschungsgemeinschaft (DFG, German Research Foundation) for funding this research: Project number 233630050-TRR 146 and 464588647-CRC 1551. AP and PV gratefully acknowledge the computing time granted on the supercomputer MOGON II and III at Johannes Gutenberg University Mainz as part of NHR South-West.

## Author contributions

BP, EK conceptualized and developed the investigation with the input from CB, TK. BP, EK, DT performed single molecule experiments. BP, EK, MDB, DT analyzed the experimental data. AP developed the simulation model and performed molecular dynamic simulations under the guidance of PV. TK, GF, EC, CC, AD purified protein complexes. EK, CB, PV, LA, KDC, HS supervised the project. EK, BP wrote the original draft. All the authors reviewed and edited the manuscript.

## Competing interests

All authors declare that they have no competing interests.

## Materials & Correspondence

Original imaging data and protein expression constructs are available upon request.

## Data availability

Source data files will be available before publication. Original imaging data are available upon request.

## Code availability

The Python-based data analysis source code used for the analysis of the imaging data will be available before publication. Simulation scripts are available upon request.

## Methods

### Protein expression and purification

#### The Smc5/6 complex overexpression and purification from S. cerevisiae

*S. cerevisiae* Hexameric (Smc5, Smc6 and Nse1-4) and octameric (Smc5, Smc6 and Nse1-6) Smc5/6 complexes were purified according to the published protocol (Pradhan et al. Nature). Briefly, the Smc5/6 complex were overexpressed by galactose-inducible system in YEP-lactate medium and isolated with tandem affinity purification system using IgG Sepharose 6 FF (VWR, 17-0969-01) and calmodulin Sepharose 4B (Merck, GE17-0529-01). The eluate was concentrated using Vivaspin 20 100K MWCO ultrafiltration unit (Sartorius, VS2041), with simultaneous exchange to STO500 buffer (50 mM Tris-HCl pH 8.0, 500 mM NaCl, 2 mM MgCl_2_, 0.5 mM tris(2-carboxyethyl)phosphine (TCEP), 10% glycerol, 0.1% IGEPAL CA-630). Fluorescently labelled hexameric Smc5/6 complexes that carry a C-terminal SNAP-tag on the Nse2 subunit or the Nse4 subunit were overexpressed, purified, and labelled according to the published protocol (Pradhan et al. Nature). Briefly, the eluate from IgG Sepharose 6 FF was incubated with SNAP-Surface-Alexa Flour 647 (New England Biolabs, S9136S) at 41⍰ºC overnight. After the reaction, the excess label was eliminated by 100K MWCO Amicon Ultra centrifugal filter (Merck, UFC5100) with concomitant buffer exchange to STO500 buffer.

#### The Smc5/6 complex overexpression and purification from E. coli

*S. cerevisiae* hexameric Smc5/6 complex (Smc5, Smc6, and Nse1-4) was also produced in *Escherichia coli* using the vectors and protocols from the Prof. S. Gruber group (Taschner et al, EMBO, 2021; Roman et al., bioRxiv 2023) with modifications. Plasmids were transformed into Rosetta 2(DE3) cells (Novagen). Liquid cultures of 2×1.5 L were prepared in Terrific Broth with antibiotics and grown in Schott flasks in a LEX bioreactor (Epiphyte3, Canada) at 37ºC until the cultures reached an optical density of approximately 2.0, measured at 600 nm. Then, the temperature was lowered to 18ºC, and protein expression was induced using 0.5 μM isopropyl ß-D-1-thiogalactopyranoside (IPTG) for 16 h. Cells were lysed by freeze-thaw cycles and sonication in lysis buffer (50 mM Tris pH 7.5, 300 mM NaCl, 5% glycerol, 2 mM TCEP, 0.01% NP40) supplemented with 2.5 kU of nuclease (Thermo Scientific, 88701) and protease inhibitor cocktail (Sigma-Aldrich, S8830). Cleared lysate was applied to a 5 ml Strep-Tactin XT 4 Flow high capacity column (IBA, 2-5028-001). After washing with 10 CV lysis buffer, bound proteins were eluted using BXT buffer (IBA, 2-1042-025). Fractions containing target protein were applied to a 5 ml HiTrap Heparin HP column (Cytiva). After washing with 5 CV low-salt HP buffer (20 mM Tris pH 7.5, 200 mM NaCl, 2 mM TCEP, 0.01% NP40), bound proteins were eluted with a 5 CV continuous gradient of 200 mM to 1 M NaCl in low-salt HP buffer. The peak fractions were concentrated using a 30 kDa MWCO Vivaspin-20 concentrator (Sartorius, VS2021) and further purified by size exclusion chromatography (SEC) using a Superdex 200 Increase 10/300 GL column (Cytiva) in SEC buffer (20 mM Tris-HCl pH 7.5, 250 mM NaCl, 1 mM TCEP and 0.05% NP40). The peak fractions were concentrated again and snap-frozen in liquid nitrogen. For purification of Smc5/6 complex with a C-terminal Halo Tag on the Nse4 subunit for fluorescent labeling, 3x StrepTrap HP 1mL columns (Cytiva) were used and proteins were eluted with lysis buffer containing 2.5 mM desthiobiotin. The SEC peak fractions were incubated with 14 µM Janelia Fluor646 HaloTag ligand (Promega, GA1120) at room temperature for 1 h and excess label was eliminated by 100K MWCO Amicon Ultra centrifugal filter with concomitant buffer exchange to STO500 buffer.

#### Wadjet overexpression and purification

The maleimide-labelable *Pseudomonas aeruginosa* PA14 JetABC complex was purified as previously described (*8*). Briefly, the His_6_-JetA + untagged JetB coexpression construct (*15*) was modified to remove native cysteine residues in JetA (C36A, C355A) and to insert a cysteine into a disordered loop region of JetA (C66; this residue was added between JetA residues A65 and S66 in the wild-type sequence). For JetC expression, a tagless JetC construct was used, as tagless JetC efficiently binds to Ni^2+^ affinity resin (*15*).

JetAB and JetC proteins were expressed in *E. coli* Rosetta2 pLysS (EMD Millipore) by growing cells in 2XYT media at 37°C until the OD_600_ reached 0.55–0.75, followed by induction with 0.33 mM IPTG. Cultures were incubated overnight (∼16 hours) at 20°C for protein expression. Cells were harvested by centrifugation, and pellets were resuspended in ice-cold buffer containing 50 mM Tris pH 7.5, 300 mM NaCl, 10 mM imidazole, 10% glycerol, and 2 mM β-mercaptoethanol. The resuspended cells were lysed by sonication, and the lysate was clarified by centrifugation. Proteins were purified using Ni^2+^ affinity chromatography (Ni-NTA Superflow, Qiagen). JetAB and JetC proteins were further purified by anion-exchange chromatography (HiTrap Q HP, Cytiva) using a buffer containing 20 mM Tris pH 7.5, 2 mM β-mercaptoethanol, and 50 mM to 1 M NaCl. The eluted proteins were concentrated and passed through a Superose 6 Increase 10/300 GL size exclusion column (Cytiva) in a buffer containing 20 mM HEPES pH 7.0, 150 mM KCl, and 1 mM TCEP (tris(2-carboxyethyl)phosphine).

For JetAB labeling with a maleimide-based fluorescent tag, the JetAB complex was mixed with Janelia Fluor 646 dye (Tocris) at a 1:20 molar ratio of JetAB to dye. The mixture was incubated overnight at 4°C with constant gentle rotation. The next day, excess dye was removed by passing the complex through a Superose 6 Increase 10/300 GL size exclusion column (Cytiva), and the conjugation efficiency was measured using a NanoDrop (Thermo Scientific). The average labeling efficiency was determined to be ∼70% for the two labeling sites in the JetA_2_B_4_ complex.

Purified JetAB (labeled JetAB) and JetC subunit proteins were mixed in a specific stoichiometric ratio to obtain the JetA_2_B_4_C_4_ complex. Any aggregated particles were removed through another round of size exclusion chromatography. The purity of the samples was assessed using SDS-PAGE analysis, and the samples were flash-frozen in liquid nitrogen and stored at -80°C until use.

#### Cohesin overexpression and purification

Human cohesin was purified as subcomplexes (Smc1/Smc3/Rad21 as a trimeric complex, STAG1 and NIPBL both independently) and reconstituted before use. The core Smc1-Smc3-Rad21 trimer used C-terminal tags on tag SMC3 (HRV3C-His10) and RAD21 (HRV3C-ybbR-TEV-2xStrepII). STAG1 was engineered with a C-terminal HRV3C-ybbR-TEV-2xStrepII tag. For ease of purification, a truncated variant of NIPBL (with an N-terminal deletion until P1162) was used and included an N-terminal fusion to MBP and a C-terminal HRV3C-ybbR-TEV-StrepII tag. All constructs were cloned into pLIB vectors.

These constructs were transformed into *E. coli* DH10emBacY (MultiBac) cells. The resulting transposed bacmid was isolated and used to transfect *S. frugiperda* Sf9 cells (2 mL of 0.5×10^6^ cells/mL) in Sf-900 III SFM media (Thermo Fisher Scientific) supplemented with Cellfectin II (Gibco). First-passage viruses (P1) were harvested after 72 h. These viruses were further amplified (P2) in Sf9 cells (25 mL of 0.5×10^6^ cells/mL) grown for 72 h at 27 °C with 120 rpm shaking using 2 mL of P1. P2 virus was harvested by centrifugation (1000 ×g, 22 °C) and stored at 4 °C with 2% (v/v) FBS (Gibco). Expression used 1 mL P2 per 100 mL of cells at a density of ∼2×10^6^ cells/mL. Cells were maintained at 120 rpm at 27 °C for 48-72 h and harvested by centrifugation once viability reached ∼90%. The resulting cell pellets were washed with PBS, snap-frozen, and stored at -80 °C.

Purification of the Smc1/Smc3/Rad21 trimer was completed as described previously for the tetrameric cohesin complex(*16*) with the following modifications: use of a 5 mL StrepTrap XT column (Cytiva) in place of a StrepTrap HP column (and subsequent elution with 50 mM biotin (Sigma)), cleavage of the Twin-StrepII tag off Rad21 prior to size exclusion chromatography (SEC) using AcTEV protease (16 h, 4 °C; Thermo Fisher Scientific) and substitution of ATTO647N for Alexa647 in the fluorophore conjugation step. Any residual uncleaved protein was removed from the sample by incubation with 100 µL Strep-Tactin Superflow Plus (Qiagen) for 30 min at 4 °C prior to SEC. Cohesin trimer concentration and labelling efficiency (typically 50-75%) was calculated by spectrophotometry and used a vendor-supplied correction factor of 0.03 for Alexa647 absorbance at 280 nm. Proteins were snap frozen as aliquots for storage at -80 °C.

Purification of MBP-NIPBL was completed as previously described(*16*) and purification of STAG1 utilized the same methodology.

#### Condensin overexpression and purification

The pentameric *S. cerevisiae* condensin complex was prepared according to previously published expression and purification protocols (*1*).

### Single-molecule imaging of DNA loop extrusion by SMC complexes

Loop extrusion assay was performed as described in Pradhan et al. (*6*) Briefly, λ-DNA (unless otherwise specified) with biotin at both terminals was anchored to glass slides functionalized with streptavidin in a flow cell channel. DNA solution was introduced at a flow rate of 2-3 μl min^−1^, resulting in an average end-to-end distance of approximately 5 μm..

Loop extrusion was initiated by introducing SMC complexes in imaging buffer. Unless otherwise indicated, the imaging buffer consisted of 100 nM SYTOX Orange, 100 mM NaCl, 7.5 mM MgCl_2_ in TX-buffer (100 mM Tris-HCl pH 7.5, 0.5 mg ml^−1^ BSA, 0.2 mM TCEP, 1 mM ATP, 30 mM d-glucose, 2 mM trolox, 10 nM catalase, 37.5 μM glucose oxidase). Specific salt concentrations, temperatures, and DNA substrates for different SMC complexes are described below.

Typically, 10,000 images with an acquisition time of 100 ms (unless otherwise specified) were recorded and stored for subsequent analysis. For wild-type non-labeled SMC complexes, solely a 561 nm laser was used to visualize the DNA. In the case of fluorophore-labeled SMC complexes, both 561 nm and 640 nm lasers were irradiated alternately to image both the DNA and the SMC complexes.

#### Smc5/6

0.5-1 nM WT Smc5/6 was utilized for loop extrusion in an imaging buffer containing 100 mM NaCl, 7.5 mM MgCl_2_, and 100 – 200 nM SYTOX Orange at 30°C. For fluorophore-labeled Smc5/6 complexes, concentrations of 1-2 nM were used.

#### Wadjet

A 44 kbp DNA substrate was utilized for loop extrusion with Wadjet as previously described (*17*). A 100 pM of fluorophore-labeled JetABC in an imaging buffer containing 100 mM NaCl, 10 mM MgCl_2_, and 100 nM SYTOX Orange was introduced to the flow cell at 30°C.

#### Cohesin

For determination of the loop extrusion directionality by cohesin (Fig 2 n,o,p), the dependency of the ratio of rates on DNA tension (Fig 4 d) and single-tethered DNA experiments (Fig S10 k, n) cohesin purified as tetramer was used as follows: Cohesin trimer, STAG1 and NIPBL were mixed at a molar ratio of 1:2.5:2.5 and incubated for 1 min at room temperature prior to dilution to the final concentrations. in imaging buffer containing pH 7.4 50 mM Kglu 2.5 mM MgCl_2_ and 200 nM Sytox Orange at 37°C. . For single tethered DNA with buffer-flow experiments, cohesin was used at 0.5–1 nM together with a 2.5-fold molar excess of NIPBL in imaging buffer containing pH 7.4 50 mM NaCl 7.5 mM MgCl_2_ and 200 nM Sytox Orange at 37°C. _2_.

For determination of the cohesin stoichiometry (Fig 3d, e, f), the comparison of monomer- and dimer-enriched SEC peaks (Fig 3h), the determination of loop extrusion directionality for the respective SEC peaks (Fig 3i) and for side flow experiments with double tethered DNA and buffer flow (Fig S10a) cohesin purified as trimer was used as follows. Cohesin trimer (either labeled with Alexa Fluor 647 or unlabeled) and STAG1 were mixed at a molar ratio of 1:2.5 and incubated at room temperature for 15 minutes. Subsequently, NIPBL was added at a 2.5x molar ratio to cohesin trimer and all components were incubated at room temperature for an additional 1 minute prior to dilution to the final concentrations in imaging buffer containing pH 7.4 100 mM Kglu 5 mM MgCl_2_ and 100 nM Sytox Orange at 37°C. The 44 kbp DNA substrate biotinylated at both ends was used.

#### Condensin

1 nM WT condensin was introduced in an imaging buffer containing 50 mM NaCl, 2.5 mM MgCl2, and 200 to 500 nM SYTOX Orange at room temperature.

### Single-molecule imaging analysis for loop extrusion directionality

DNA molecules were visualized using TIRF microscopy and recorded as image sequences. Individual molecules were cropped and saved as TIFF image stacks. Kinetic information, including loop positions and segment intensities, was extracted using custom software ((*18*), code available at: https://github.com/biswajitSM/LEADS).

Briefly, kymographs were constructed by summing pixel intensities across the DNA axis at each time point. Loop positions were identified, and intensities within a 9-pixel window around the loop peak were recorded. Intensities on both sides of the loop (I and II) were also collected. Segment intensities were converted to kilobases, assuming the total intensity represented the full DNA substrate length (48.5 kb for λ-DNA).

Segment sizes were smoothed using a Savitzky-Golay filter (*19*) to calculate rates of change. Rates of loop extrusion were calculated at each time point using a 5-second (⊿*t*) interval:

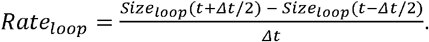

Similarly, *Rate*^I^ and *Rate*^*II*^ were calculated by using the sizes of segments I and II (Fig. 1d).

The ratio-of-rates was calculated as:

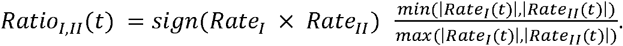

The initial growth phase was defined as the first 20 seconds during which Rate_loop_. Remained positive. Subsequent loop size changes were categorized as “regrowth” (Rate_loop_ > 0.05 kbps) or “shrinkage” (Rate_loop_ < -0.05 kbps), collectively termed the “mature phase.” Time points with Rate_loop_ between -0.05 and 0.05 kbps were excluded from further analysis.

For symmetry classification, Ratio-of-rate values > 0.1 were classified as two-sided extrusion, values < - 0.1 as one-sided with slippage, and values between -0.1 and 0.1 as one-sided extrusion (Fig. 1f, g, Fig. S1).

To ensure reliable directional assignments, time points for which |Rate_I_| or |Rate_II_| fell below 0.05 kbp/s were additionally excluded from ratio-of-rates and Rate_I_-versus-Rate_II_ analyses. This threshold corresponds to approximately half of the frame-to-frame standard deviation of DNA segment size fluctuations (0.12 kbp/s), thereby restricting the analysis to time points with a robust directional signal.

The accuracy and robustness of this rate-extraction and directionality classification pipeline were validated using two independent approaches. First, the finite difference method was benchmarked against a derivative-based approach applied to the traces (Savitzky–Golay filter, second-order polynomial, 5-second window), yielding nearly identical rate profiles and symmetry classifications (Fig. S11a–c). Second, the full analysis pipeline was applied to molecular dynamics simulation with known ground-truth extrusion rates, demonstrating that loop detection, rate extraction, and directionality classification accurately recover the input symmetry distributions for both two-sided and one-sided extruder models (Fig. S11d–m).

### Stalling force estimation

DNA tension was estimated from the extension of the DNA segments outside the loop using the worm-like chain model. At each time point, the relative DNA extension was calculated

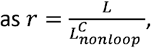

where L is the end-to-end distance between the two of DNA tethering points and 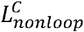 is the contour length of the DNA out the loop, calculated from the combined lengths of segments I and II and converted to micrometers.

DNA tension was then calculated as 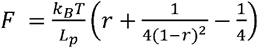, where k_B_T=4.1 pN.nm and L_p_ is the persistence length of the DNA (*20*). Both persistence length (35 nm) and DNA length per base pair (0.338nm338nm338nm338 nm338nm/bp) were taken from Davidson et al.(*21*), who measured these parameters under conditions matching those used in this study.

The stalling force of an extrusion event was estimated from the DNA tension at maximal loop size. Specifically, we identified time points at which the loop size exceeded 95% of the maximum loop size observed at least 50 s after loop initiation and calculated the mean DNA tension across these time points.

To obtain the relationship between tension and the ratio-of-rates (Fig. 5a-d), logarithmically spaced bins were created, spanning the full range of tensions observed for a given SMC complex across all molecules. Ratio-of-rates values were assigned to the corresponding tension bins, and the mean and standard deviation were calculated for each bin.

### Photobleaching stoichiometry analysis

Fluorescence intensity traces were extracted from pixels corresponding to the position of the loop anchor point in each kymograph. The number of discrete photobleaching steps in each trace was counted manually to determine the number of fluorophore-labeled subunits present per complex. For events in which no photobleaching occurred within the observation period either because the fluorophore had already bleached before the event began or because bleaching was not observed; the fluorophore count was estimated by comparison to the fluorescence intensity of either a previously bleached molecule on the same DNA molecule or a surface-immobilized reference molecule in the same field of view. Events were assigned to one of four categories: 0, 1, 2, or >2 fluorophores detected.

To assess whether the observed fluorophore count distribution is consistent with a monomeric or dimeric complex, we constructed theoretical probability distributions for each model based on the experimentally determined degree of labeling f (i.e., the fraction of subunits carrying a fluorophore). For a dimeric complex in which each subunit is independently labeled with probability f, the expected probabilities are:

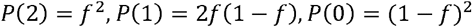

For a monomeric complex carrying a single labelable subunit:

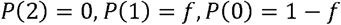

The degree of labeling f for each complex was determined independently by absorbance measurements (see Protein purification and labeling). The theoretical distributions were then overlaid on the experimental fluorophore count histograms to assess goodness of fit visually and quantitatively (Fig. 3a– c; Fig. S5).

For the stoichiometry analysis of surface-immobilized cohesin complexes (Fig. S5h-i), fluorescence intensity traces were compiled into a histogram that exhibited two distinct populations corresponding to one and two fluorophores. The mean intensity of the single fluorophore population was used to calibrate the two-fluorophore population, and the relative areas of the two peaks provided were used to estimate the monomeric and dimeric fractions.

### MSD analysis

To validate the time window used for rate calculations, we quantified the positional fluctuations of DNA segments and loop positions using mean-square displacement (MSD) analysis. For each trajectory, position coordinates were extracted frame-wise with the peak finding algorithm, and a representative particle track was mean-centered prior to analysis. MSD was then computed with trackpy.imsd. The TrackPy input table included frame, particle, x, and y; because y was fixed to a constant value, the reported MSD corresponds effectively to one-dimensional fluctuations along the x axis. Results were plotted as MSD versus lag time on logarithmic axes (Fig. S2l,m), and used to verify that the 5-s interval employed for rate calculations was sufficiently long to suppress positional noise while remaining short relative to the timescale of loop extrusion dynamics.

### Switching direction

Direction switching events during one-sided extrusion were identified automatically. The Rate_I_ and Rate_II_ values were smoothed using a Savitzky-Golay filter with a window length of 5 seconds. Time points where either Rate_I_ or Rate_II_ crossed zero were initially identified as potential crossing points. To qualify as a true direction-switching point, the crossing point had to meet the following criteria: 1) The mean values of both Rate_I_ and Rate_II_ since the last crossing point exceeded a threshold of 0.05 kbps. 2) After the crossing point, Rate_I_ and Rate_II_ had opposite signs. 3) Both Rate_I_ and Rate_II_ had changed its sign since the last switching point. The number of switching events per molecule was normalized to 100 seconds of observation time. Mean and standard deviation of switching frequencies were then calculated across all molecules (Fig. 5g).

### Fluorescent labeling efficiency estimation for E.coli expressed Smc5/6

The concentration of E.coli-expressed Smc5/6 labeled with JF646 was estimated using a Bradford assay with BSA as the standard. This sample was then diluted to a 1 μM stock concentration. Subsequently, the concentration of JF646 was determined through fluorescence correlation spectroscopy (FCS), using a Zeiss LSM 880 Observer Airyscan microscope. 1 nM ATTO647N was used to calibrate for the detection volume of the confocal microscope. For FCS analysis, the stock Smc5/6 sample was further diluted to 1 nM in the imaging buffer, and autocorrelation data was recorded. The resulting autocorrelation curves were fitted with both diffusion and blinking components, in the Zeiss software. This analysis resulted in a JF646 concentration of 0.65 nM, implying a 65% labeling efficiency. However, a subsequent observation was made: post-incubation of the sample in an ice bath for an hour, the labeling efficiency decreased to 40%. Thus, we establish a labeling efficiency range of 40% to 65% for the *E. coli*-expressed Smc5/6.

### Molecular Dynamic Simulation

Coarse-grained simulations of DNA are based on a standard bead-spring model: Beads are connected with FENE springs and chain stiffness is implemented with a cosine-type potential with amplitude *κ*. Monomers interact via a repulsive Weeks-Chandler-Andersen potential which accounts for excluded volume effects. Simulation parameters (unit of length *σ*, numbers of beads in a chain N and stiffness *κ*) are matched with experiments via an approach detailed in Ref. (*22*): As charges are not modelled explicitly, we determine the effective diameter with an approach based on polyelectrolyte theory for given ionic conditions. In addition, we match persistence and contour length, which yield *κ* and N respectively. This model was shown to reproduce structural and topological properties of DNA in good agreement with corresponding experiments (*23, 24*). All parameters used in this work are given in table 1.

**Table 1:** Simulation parameters representing *λ*-DNA (48502 bp), ionic conditions equivalent to 115mM NaCl and a persistence length of 35nm.

|  |  |
| --- | --- |
| Number of monomers | 3026 |
| Default graft distance | 5μm |
| FENE constant K | 30 $\epsilon$/ $\sigma^2$ |
| FENE constant $r_0$ | 1.5 $\sigma$ |
| Bending $\kappa$ | 6.549 $k_B T$/ $\sigma$ |
| WCA constant $\sigma$ | 1 |
| WCA constant $\epsilon$ | 1 |
| Friction constant $\gamma$ | 0.1 $\tau^{-1}$ |

The SMC protein complex is modeled using a simplified handcuff structure, consisting of two rings, each composed of 10 monomers as proposed in Ref. (*25*). By default, this handcuff structure is initially placed at the center of the DNA polymer chain and the chain is allowed to equilibrate. These rings act as openings through which the polymer is extruded or slipped during the simulation. This double loop structure (*25, 26*) has the advantage of simpler representations such as dynamic bonds between opposing sites of DNA (*27, 28*) that it allows for the implementation of an explicit driving force, which is given in units of k_B_T/σ, and a more detailed distinction between two-sided and one-sided extruders. The polymer’s ends are tethered to a wall which is interacting with the polymer via a 9-3 Lennard-Jones potential, which is obtained by integrating the standard Lennard-Jones potential over a semi-infinite continuum wall. The mass of all beads is fixed to 1, while the mass of the center of mass of the handcuff is set to 5 (*25*). To perform extrusion, we compute the component normal to the plane of each of the rings and give an extrusion force to the monomers which are closest to the respective rings along these normal components. Corresponding equal and opposite forces are applied to the handcuff for equivalence. Using this approach, we can simulate one-sided or two-sided extrusion by varying the magnitude of force each of the ring applies.

Note that for the computation of stalling and extrusion forces in experimental units, we apply the Marko-Siggia equation (see section on stalling forces), which is based on the extension of the unextruded fraction of DNA. Dimeric extruder simulations, for instance, which require a force of 0.08 k_B_T/σ or 0.06pN (corresponding to Smc5/6) give rise to an extension that correspond to a stalling force of about 0.2 pN. Note that there is no one-to-one correspondence between forces applied to the rings and stalling tensions due ring geometry and entropic contribution of the loop.

We apply a small force (0.013 k_B_T/ σ) to the second ring during one-sided extrusion to mimic anchoring of one-sided motors. The magnitude of this force is such that it cannot cause any extrusion on its own and just opposes the slipping that happens during one-sided extrusion. This provides a more realistic representation of slippage than just fixing the bond.

While our coarse-grained model already provides a mapping of simulation to experimental length scales, i.e., one bead roughly corresponds to 16 base pairs, the matching of time scales is more involved. There are two time scales of interest in the system: (1) the diffusive time scale that can be obtained by comparing the mean square displacement of the central segment with corresponding experiments and; (2) the faster extrusion time scale, obtained by matching the initial extrusion rates between experiments and simulations, which is used throughout this work to speed up simulations. For Smc5/6, 0.5 x 10^6^ time steps=0.5 x 10^4^*τ* in simulations roughly correspond to one second in experiments, and for convenience, we used ∼0.5 x 10^6^ time steps as one second for analysis of simulations. Note that the two time scales could be matched by introducing waiting times during which the DNA is attached to the handcuff and extrusion does not proceed.

For simulations we use the molecular dynamics package HOOMD-blue (*29*) on GPUs and CPUs. Equations of motion are integrated using a standard Langevin thermostat with dt = 0.01τ and *γ*= 0.1. We compared simulations across different values of γ, demonstrating that it functions as a temporal scaling parameter. The resulting diffusion curves can be rescaled to overlap with the experimental curve (*12*), enabling substantial computational acceleration while preserving the dynamics of interest. When calculating rates (loop, I, II), a 5-second interval (equivalent to ∼2.5 x 10^6^ time steps) was employed. For ratio-of-rates and RateI vs. RateII analyses, time points where |Rate_I_| or |Rate_II_| fell below 0.03 kbp/s were excluded. This threshold was set to approximately half the frame-to-frame standard deviation of DNA segment size changes in simulations (0.06 kbp/s), consistent with the exclusion criterion applied to experimental data.

For detailed discussions of time scale matchings, simulation parameters, the applicability of the Marko-Siggia equation and mappings of SMC5/6, Wadjet and cohesion to the handcuff model, the reader is referred to Pinto et al (*12*).

### Modeling Loop Extrusion Symmetry from Experimentally Observed Translocation Rates

We obtained translocation speeds for each Smc5/6 molecule by measuring their speed within three-second intervals, recording these along with the end-to-end distance of the DNA they translocated on (Fig. S8a-f). Analyzing DNA with different end-to-end distances (representing various relative extensions) provided translocation speeds at different relative extensions. Speed values were considered positive, as the initial direction for each molecule is random.

To model two physically linked translocating motors, we randomly selected two translocation speeds (for Rate_I_ and Rate_II_) from DNA molecules with the same relative extension. These rates were assigned negative signs, reflecting their action of reeling in DNA from outside to inside the loop, thus decreasing the sizes of segments I and II outside the loop. A 2D histogram was generated by extracting 10,000 such pairs of translocation speeds (Fig. S8g). However, slippage with positive segment rates was not modeled as translocation data lacks slippage information, resulting in population exclusively in Q3. This approach models loop extrusion rate variation without considering time/phase-dependent DNA tension changes.

## Notes

### Competing Interest Statement

The authors have declared no competing interest.

### Summary of Updates

Summary of major changes: 1. New photobleaching and SEC data establish that condensin is monomeric, while cohesin, Smc5/6, and Wadjet are dimeric. 2. Added extensive controls, including high-resolution tracking, tagging controls, and MD simulation calibrations. 3. Mechanistic reframing: We now show one-sided extrusion and direction switching emerge from stochastic motor competition near the stalling regime, rather than active switching states. 4. New simulations (Figs. 4-5) demonstrate that increasing DNA tension destabilizes two-sided extrusion in dimeric motors, explaining cohesin's instability. 5. Expanded Discussion provides a unified physical framework and reconciles previous conflicting reports (e.g., Barth et al.). 6. Revised title and manuscript narrative to reflect the central role of DNA tension and motor stoichiometry.

