## Supplementary Information for "A DNA tension-dependent tug-of-war between dimeric SMC motors governs loop extrusion directionality"

**The file includes:**

Supplementary Text

Figs. S1 to S11

**Supplementary Note 1: Experimental Factors Influencing Loop Extrusion Directionality**

Because loop extrusion directionality reflects the mechanical state of the extruding complex, it is expected to respond sensitively to any perturbation that alters DNA tension or motor coordination. Consistent with this expectation, we find that directionality is influenced by several physicochemical parameters across our experiments.

Protein tagging. Fluorescent tagging of Smc5/6 at the Nse4 or Nse2 subunit shifted extrusion behavior toward more one-sided, slippage-dominated states relative to untagged wild-type complexes, increasing loop mobility approximately 3–4 fold (Fig. S4a–h). Ratio-of-rates distributions for tagged proteins were skewed toward negative values in all extrusion phases (Fig. S4i–n), indicating that tagging modestly perturbs the mechanical stability of the extruding complex. Importantly, however, directionality fractions remained unchanged across fluorophore occupancy classes within each tagged complex (Fig. S4l–n), confirming that the tag—rather than the fluorophore itself—is the source of the perturbation. These observations highlight the importance of considering tagging strategies when comparing extrusion directionality measurements across studies.

Buffer flow (side-flow). Real-time visualization of loop extrusion typically involves applying constant buffer flow at a large angle (~90°) relative to the double-tethered DNA. To investigate the effect of such forces, we simulated dimeric loop extrusion in the presence of a perpendicular drag force acting on the DNA polymer (Fig. S10b-h). Increasing flow promoted two-sided extrusion, shifting data points toward the Q3 diagonal and ratio-of-rates values toward unity (Fig. S10b–f). This enhancement likely arises because the drag force counteracts DNA slippage from the passive side of the loop. Consistent with this interpretation, externally applied side-flow promoted two-sided extrusion by dimeric cohesin (Fig. S10b-h) as reflected by more points along the diagonal in Q3 quadrant at higher flow rate. Together, these findings suggest that relatively small external forces can influence loop extrusion directionality and should be considered when comparing experiments performed under different flow conditions.

Ionic conditions and DNA staining. Additional control experiments confirmed that ionic conditions and SYTOX Orange DNA staining concentration influence the magnitude of observed directionality transitions but do not alter the qualitative directionality patterns for any SMC complex (data not shown). These observations further underscore that loop extrusion directionality is a mechanically sensitive property that responds to multiple features of the experimental environment.

Taken together, these results support a view in which loop extrusion directionality is not an intrinsic binary property of an SMC complex, but rather an emergent behavior that depends on the balance of forces acting on the extruding motor and the DNA substrate.

Supplementary Note 2: Loop Extrusion Directionality on Single-Tethered DNA: Two-sided extrusion by dimeric SMC complexes under minimal DNA tension

To directly test whether the progressive transition to one-sided extrusion is driven by tension accumulation during loop growth, we monitored SMC-mediated loop extrusion on single-tethered DNA, in which one end of the DNA is anchored to a surface while the other remains free (Fig. S10i-n). This configuration minimizes tension accumulation during extrusion because the free end can move as the loop grows. The primary mechanical load arises from the mild buffer flow required for visualization (~0.07 pN), which is substantially below the stalling forces of Smc5/6 and Wadjet.

Under these low-tension conditions, Smc5/6 (Fig. S10i) and Wadjet (Fig. S10j) formed loops with simultaneous shortening of DNA segments from both sides, indicating two-sided extrusion. Condensin, used at low concentrations to prevent z-loop formation by multiple complexes, extruded loops with shortening of DNA from only one side; either from the tethered end (Fig. S10l) or from the free end (Fig. S10m); demonstrating one-sided extrusion. Cohesin similarly extruded loops one-sidedly (Fig. S10k). Quantification confirmed that Smc5/6 and Wadjet exhibited near 100% two-sided extrusion, while condensin and cohesin showed 100% one-sided extrusion under these conditions (Fig. S10n).

Consistent with the reduced tension in the single-tethered geometry, loops did not enter the mature phase observed on double-tethered DNA. Instead, loops continued to grow until the extruding complex reached the free DNA end, at which point extrusion terminated or the loop disrupted abruptly. Consequently, neither the alternating shrinkage nor regrowth of the flanking DNA segments characteristic of mature-phase dynamics was observed, and frequent direction switching was absent. Together, these observations indicate that both the mature phase and the associated switching behavior arise from tension accumulation during loop growth rather than from an intrinsic transition in motor behavior.

The recovery of nearly complete two-sided extrusion by Smc5/6 and Wadjet under these conditions further demonstrates that the transition from coordinated to asymmetric extrusion is reversible and governed by DNA tension. We note, however, that the residual force generated by buffer flow is inherently directional in the single-tethered geometry, creating a persistent asymmetry between the two loop flanks. As a result, this assay is not ideally suited for probing switching dynamics independently of tension effects. In addition, the applied flow force (~0.07 pN) approaches the stalling-force regime of cohesin and may therefore contribute to its predominantly one-sided behavior in this assay.

Supplementary Note 3: Smc5/6 Single-Motor Translocation and Its Relationship to Loop Extrusion.

Our model of two-sided loop extrusion by dimeric Smc5/6 and Wadjet proposes two physically linked but mechanically independent motors, each capable of DNA translocation. This is supported by previous single-molecule observations of individual Smc5/6 complexes translocating along DNA in the presence of Nse5/6 (Pradhan et al., 2023), and by our coarse-grained simulations in which single-ring motors predominantly translocate rather than form stable loops.

To further investigate the relationship between translocation and loop extrusion, we analyzed translocation speeds of individual Smc5/6 complexes at different DNA extensions (Fig. S8b,c). This analysis revealed that translocation speed is DNA tension-dependent, mirroring the tension dependence observed for loop extrusion rates. Importantly, even at the same DNA extension, individual translocation speeds varied substantially between molecules and within single translocation events over time (Fig. S8b–f). This rate heterogeneity closely mirrors the variability in DNA reeling rates (Rate_I_ vs. Rate_II_) observed for dimeric Smc5/6 during loop extrusion (main text Fig. 2a).

To test whether independent sampling of two such translocation rates could reproduce the broad bivariate rate distributions observed during loop extrusion, we modeled two-sided extrusion by randomly pairing translocation rates from molecules measured at the same DNA extension. The resulting 2D rate distribution (Fig. S8g) was consistent with the experimentally observed quadrant plots for Smc5/6, recapitulating the broad dispersion of Rate_I_ and Rate_II_ without requiring any explicit rate-coupling mechanism. Coarse-grained simulations of a single-ring handcuff motor similarly exhibited heterogeneous translocation rates (Fig. S8h,i), confirming that rate variability is an intrinsic consequence of thermal fluctuations acting on individual motors.

Together, these results provide independent mechanistic support for the dimeric motor model: the stochastic heterogeneity in single-motor translocation rates naturally accounts for the variable and often asymmetric reeling rates observed during two-sided loop extrusion, without invoking any regulatory asymmetry between the two motors.

**Supplementary Note 4:** **Possible origins of the discrepancies between our observations of Smc5/6 loop extrusion and those reported by Barth et al.**

Our findings on Smc5/6 loop extrusion symmetry significantly differ from those of Barth et al. (*9*). We observed two-sided extrusion by dimeric Smc5/6, contrary to their conclusions. Several factors may contribute to these discrepancies:

(i) Analysis method: Barth et al. assigned a single symmetry score to each DNA loop extrusion event, representing a statistical average. Our approach, in contrast, analyzed symmetry at each time point, revealing distinct phases of loop extrusion with varying symmetries and uncovering hidden properties like the influence of tension on symmetry.

(ii) Protein expression, purification and labeling: Barth et al. used two protein expression systems (E. coli and budding yeast) and reported that E. coli expressed Smc5/6 performs loop extrusion as single complexes (with ~60% labeling efficiency). We purified both labeled and unlabeled Smc5/6 from E. coli and assessed loop formation symmetry and the number of complexes involved (Fig. S3o-s). Our photobleaching data showed a higher proportion (60%) of two-step bleaching events compared to single-step events (30%) (Fig. S3o), suggesting dimer-mediated loop extrusion (with ~70% labeling efficiency, Methods). We also observed predominantly two-sided extrusion during initial loop growth, transitioning to one-sided extrusion in the mature phase (Fig. S3p-s), similar to yeast expressed Smc5/6. Together, these observations indicate that neither the protein expression system nor fluorophore labeling alone can account for the differences between the two studies. Nevertheless, we cannot exclude the possibility that differences in purification procedures, protein quality, or assay conditions contribute to variations in the observed stoichiometry and extrusion behavior. In addition, the substantially lower labeling efficiencies reported by Barth et al. would be expected to reduce the frequency of multi-step photobleaching events and could complicate stoichiometry estimation from fluorophore counting alone.

More generally, our results demonstrate that loop extrusion directionality is highly sensitive to DNA tension and other experimental parameters (Supplementary Note 1). Consequently, differences in assay geometry, force regimes, protein preparation, and analysis methodology may all contribute to the contrasting conclusions reached by the two studies.

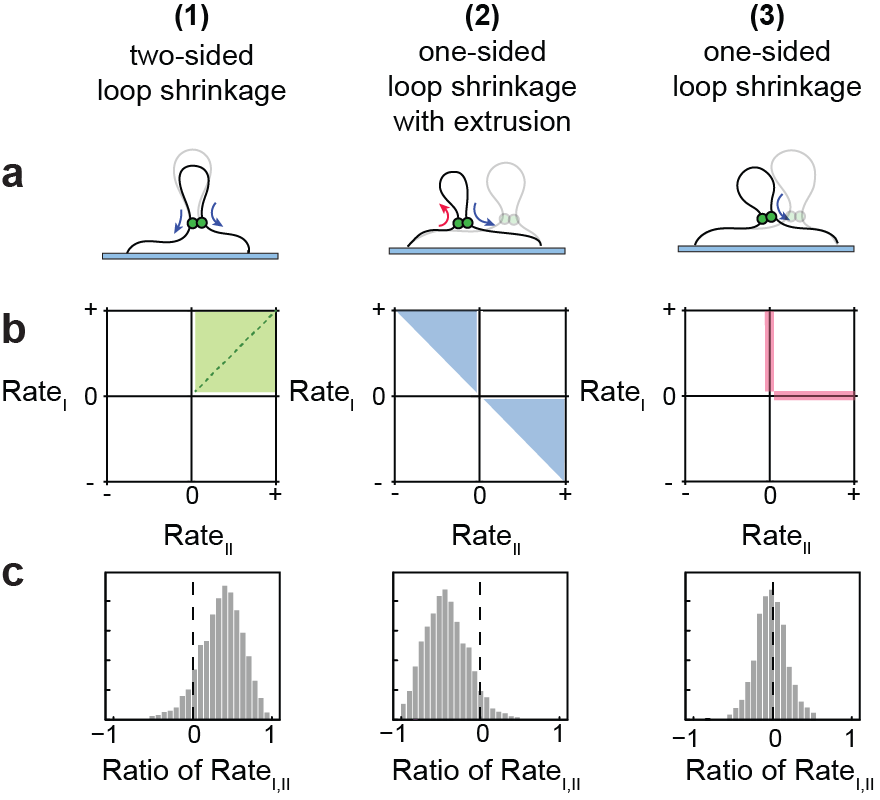

**Supplementary Figure S1. Classification of loop shrinkage symmetry.**
(a) Schematic illustrating the three symmetry classes observed during loop shrinkage: two-sided loop shrinkage (1), one-sided loop shrinkage accompanied by loop extrusion from the opposite side (2), and strictly one-sided loop shrinkage with no change in the opposite DNA segment (3). (b) RateI-versus-RateII quadrant plot, with shaded regions indicating the corresponding symmetry classes. (c) Schematic distributions of the ratio-of-rates for the three loop shrinkage symmetry classes.

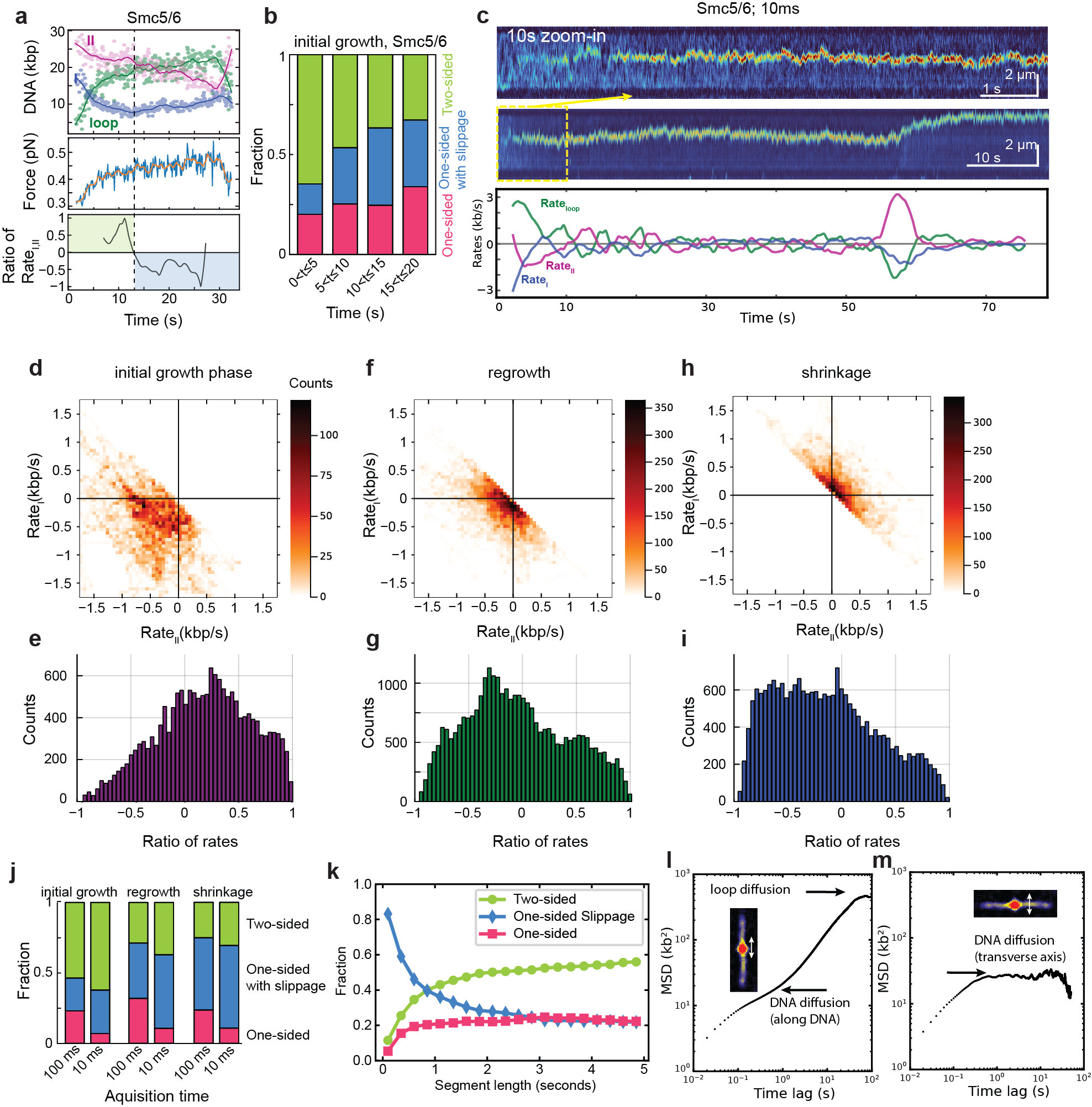

**Supplementary Figure S2. Time-dependent directionality transition of Smc5/6 loop extrusion and validation at increased temporal resolution.** (a) Representative Smc5/6 loop extrusion event showing DNA segment sizes (top; segments I, II, and loop), DNA tension (middle), and the corresponding ratio-of-rates (bottom). The ratio-of-rates transitions from values near +1 to values near 0 during loop growth. (at ~13 s), coinciding with increasing DNA tension. (b) Directionality fractions during the initial growth phase of Smc5/6 loop extrusion, binned into 5-s intervals. (c) Smc5/6 loop extrusion recorded at 10 ms temporal resolution. Top: zoomed-in kymograph showing a 10-second window (upper) and full kymograph (middle) at 10 ms acquisition time. Bottom: corresponding time traces of Rate_loop_ (pink), Rate_I_ (green), and Rate_II_ (magenta), showing that two-sided extrusion (simultaneous negative Rate_I_ and Rate_II_) is resolved at the 10 ms timescale with a 5-second window for rate calculations. (d–i) Directionality analysis of Smc5/6 loop extrusion events recorded at 10 ms temporal resolution. Quadrant plots of Rate_I_ versus Rate_II_ during initial growth (d), mature regrowth (f), and shrinkage (h), and their corresponding ratio-of-rates distributions (e, g, i). (j) Comparison of directionality fractions obtained from data acquired at 100 ms and 10 ms temporal resolution. (k) Directionality fractions as a function of the time window used for rate calculations. (l, m) Mean squared displacement (MSD) analysis of loop positions along the DNA axis (l) and perpendicular to the DNA axis (m). The MSD analysis was used to establish a lower limit of approximately 1 s for the time window used in directionality analysis, below which positional fluctuations dominate the measured rates.

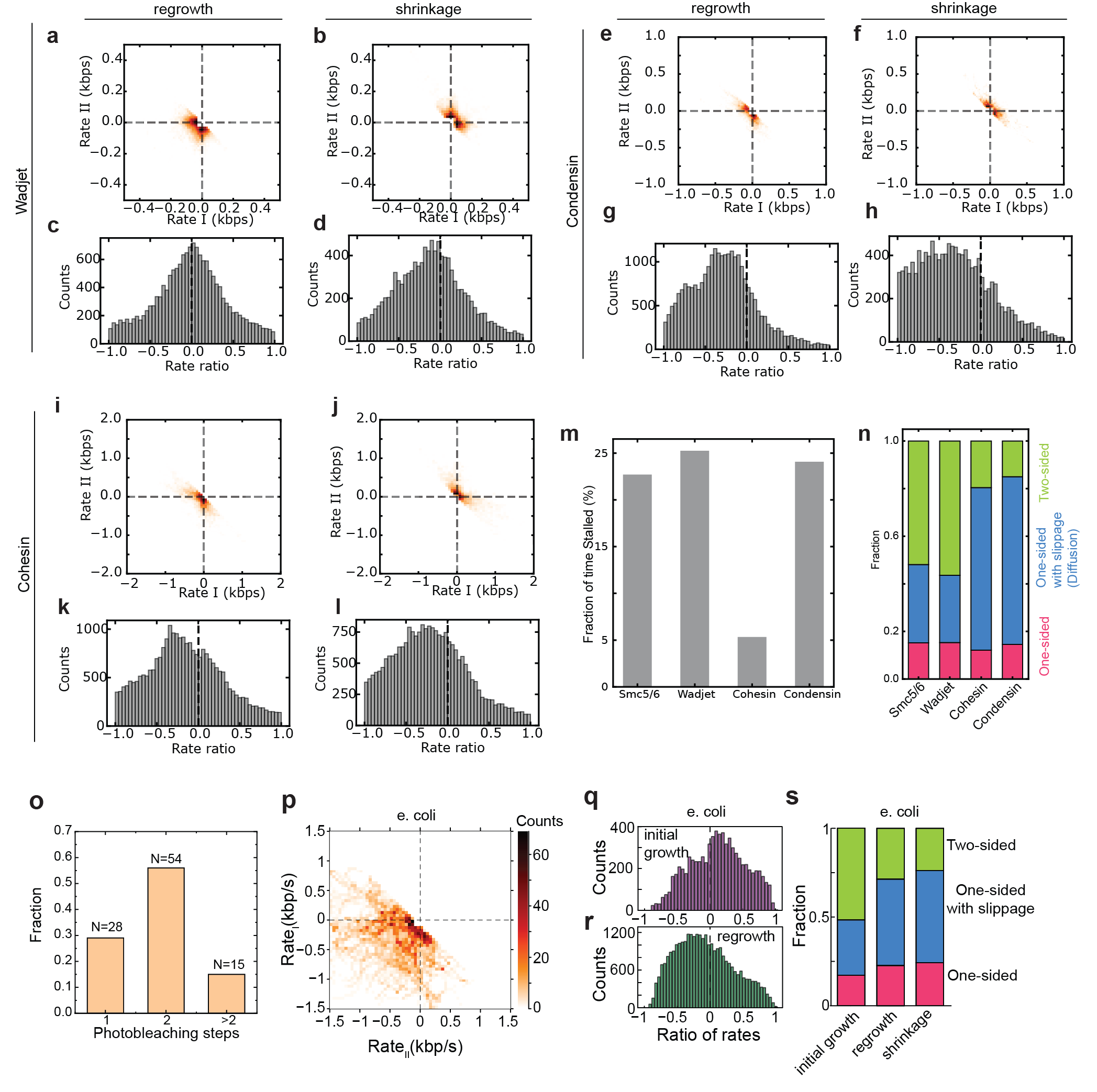

**Supplementary Figure S3. Loop regrowth and shrinkage symmetry.** (a-d) *Wadjet*: Quadrant plots of Rate_I_ vs. Rate_II_ (a, b) and histograms (c, d) of the ratio-of-rates for the mature loop regrowth (a,c) and shrinkage phase (b,d). (e-h) *Condensin*: Quadrant plots of Rate_I_ vs. Rate_II_ (e, f) and histograms (g, h) of the ratio-of-rates for the mature loop regrowth (e,g) and shrinkage phase (f,g). (i-l) *Cohesin*: Quadrant plots of Rate_I_ vs. Rate_II_ (i, j) and histograms (k, l) of the ratio-of-rates for the mature loop regrowth (I,k) and shrinkage phase (j,l). (m) Fraction of total analysis time spent stalled (both Rate_I_ and Rate_II_ within the noise band, |Rate_loop| < 0.05 kb/s) for Smc5/6, Wadjet, cohesin, and condensin. (n) Composition of the extruding (non-stalled) time for each complex, classified from the ratio-of-rates into two-sided (green; ratio > 0.1), one-sided with slippage / loop diffusion (blue; ratio < −0.1), and one-sided without slippage (pink; −0.1 < ratio < 0.1).(o-s) Symmetry and stoichiometry of E. coli expressed Smc5/6. (o) Distribution of photobleaching steps for Smc5/6 labeled with JF646 (labeling efficiency of 65%). (p) Quadrant plots showing the relationship between Rate_I_ and Rate_II_ during the initial growth phase of loop extrusion events. (c,d) Distribution of the ratio-of-rates during the initial growth (q) and mature growth phase (r) of loop extrusion events. (s) Fractions of two-sided, one-sided, and one-sided with slippage for different phases.

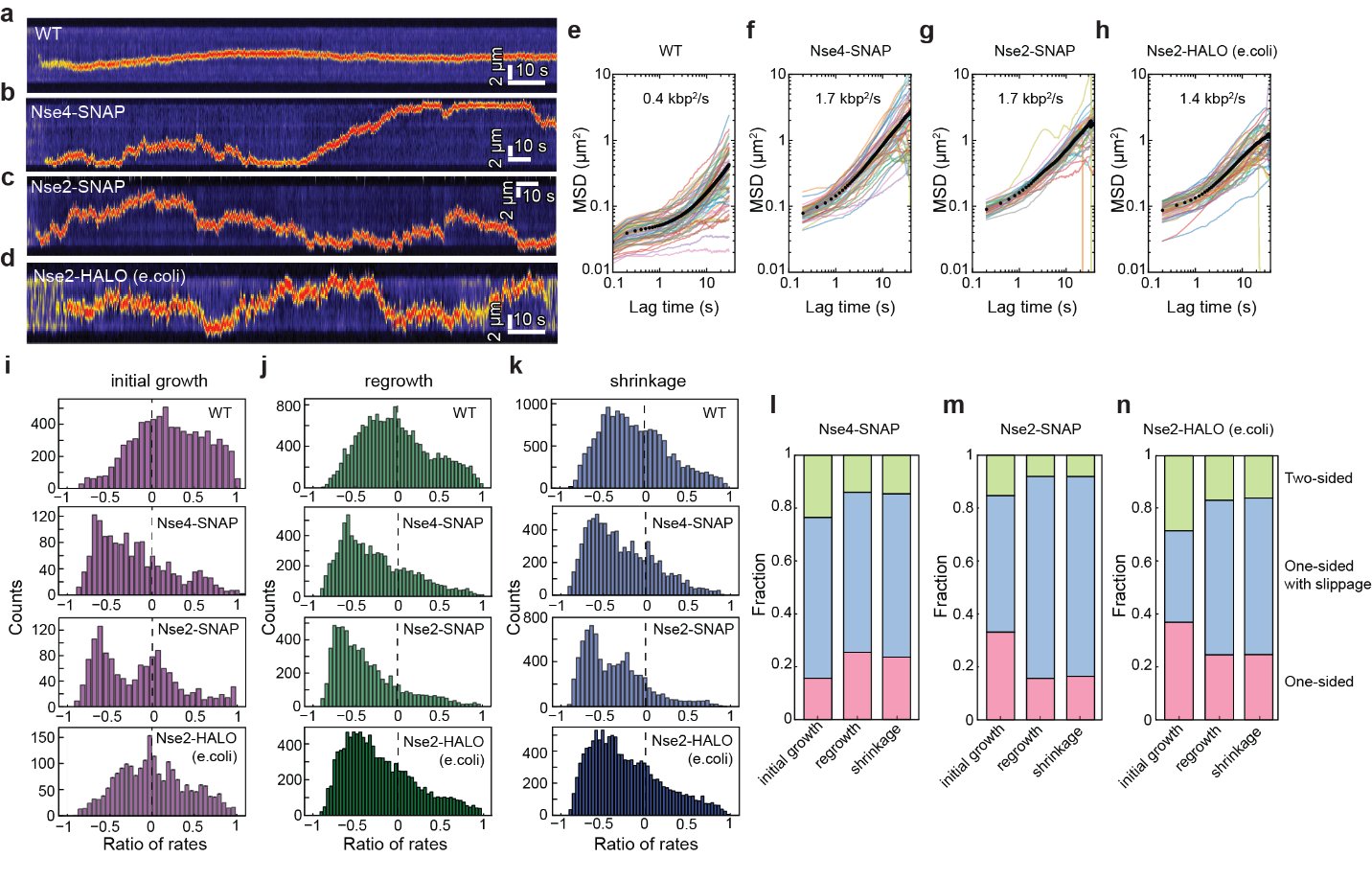

**Supplementary Figure S4. Effect of SNAP and HALO tags on loop extrusion dynamics and the symmetry evaluation.**

(a–d) Representative kymographs of loop extrusion by wild-type yeast-expressed Smc5/6 (a), SNAP-tagged Smc5/6 labeled at Nse4 (b), SNAP-tagged Smc5/6 labeled at Nse2 (c), and HALO-tagged *E. coli*-expressed Smc5/6 labeled at Nse2 (d). (e–h) Mean-square displacement (MSD) analysis of loop positions for wild-type (e), Nse4-SNAP (f), Nse2-SNAP (g), and Nse2-HALO (h) Smc5/6 complexes, showing increased loop mobility for tagged complexes relative to wild type.

(i–k) Ratio-of-rates distributions for the different Smc5/6 constructs during the initial growth (i), mature regrowth (j), and shrinkage (k) phases of loop extrusion. (l–n) Fractions of extrusion symmetry classes for Nse4-SNAP (l), Nse2-SNAP (m), and Nse2-HALO (n) Smc5/6 complexes during the corresponding extrusion phases.

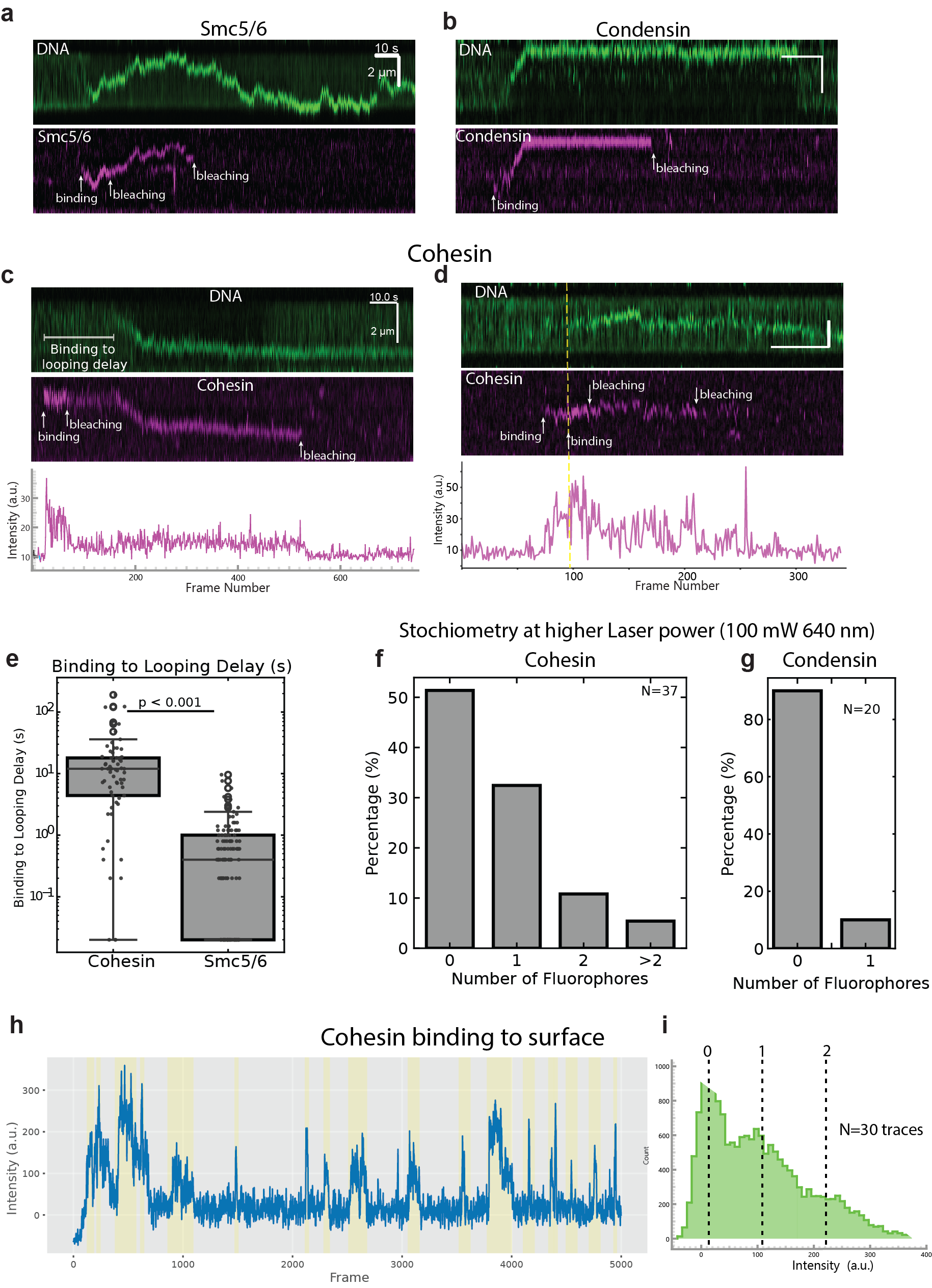

**Supplementary Figure S5. Single-molecule stoichiometry analysis of loop-extruding SMC complexes.** (a) Representative kymograph showing a DNA loop extrusion event by Alexa647-labeled Smc5/6–Nse2-Snaptag (magenta, bottom) on λ-DNA (green, top). Binding and two sequential photobleaching steps are indicated by arrows.

(b) Representative kymograph of a loop extrusion event by ATTO647N-labeled condensin (magenta, bottom) on λ-DNA (green, top), with a single binding event and single bleaching step indicated.

(c) Representative kymograph of a loop extrusion event by Alexa647-labeled cohesin (magenta, middle) on λ-DNA (green, top). A single binding event followed by two photobleaching steps is annotated. The corresponding fluorescence intensity trace of the cohesin signal over time is shown below. Note the delay between cohesin binding and loop extrusion onset ("binding to looping delay").

(d) As in (c) but showing a cohesin molecule with two sequential binding events and two bleaching steps, indicative of dimerization on the DNA. The dashed yellow line marks the time of the second binding event. The corresponding intensity trace is shown below.

(e) Box-and-scatter plot comparing the binding-to-looping delay times for cohesin and Smc5/6. Cohesin exhibits a significantly longer delay than Smc5/6 (p < 0.001, indicated by the horizontal bar).

(f, g) Bar charts showing the distribution of photobleaching step counts for DNA loop extrusion events by cohesin (f, N = 37) and condensin (g, N = 20), acquired at higher laser power (100 mW, 640 nm laser) to improve bleaching detection. The majority of cohesin events show zero or one bleaching step, while condensin events are predominantly zero-step, consistent with predominantly monomeric engagement.

(h) Representative fluorescence intensity trace of cohesin non-specifically adsorbing to the passivated glass surface, recorded over ~5,000 frames. Yellow shading highlights individual binding events used for stoichiometric analysis.

(i) Histogram of fluorescence intensities from surface-bound cohesin molecules (N = 30 traces), derived from the binding events highlighted in (h). Dashed lines indicate intensity levels corresponding to zero, one, and two fluorophores, enabling calibration of the single-fluorophore intensity and determining of labeling stoichiometry in the solution during the looping assay.

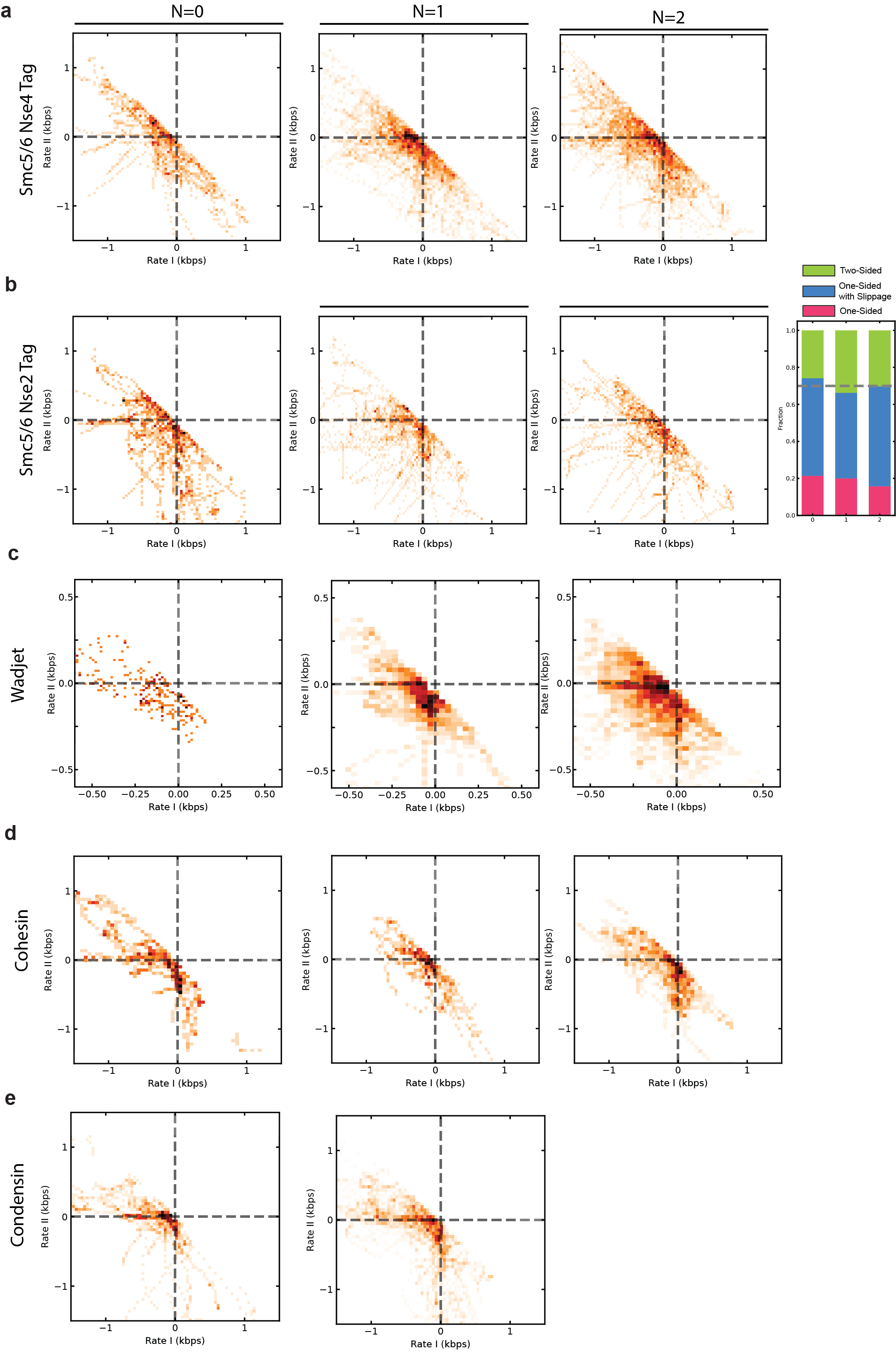

**Supplementary Figure S6. Directionality of loop extrusion is independent of fluorophore subpopulations across SMC complexes.**

Quadrant plots of RateI versus RateII for loop extrusion events during the initial growth phase, grouped by the number of fluorophores (0, 1, or 2) detected per SMC complex by photobleaching analysis. Rows correspond to Smc5/6 labeled at Nse4 via SNAP-tag (a), Smc5/6 labeled at Nse2 via SNAP-tag (b), Wadjet (c), cohesin (d), and condensin (e). Within each row, quadrant plots are shown for events containing zero (left), one (middle), or two (right) detected fluorophores. Dashed lines indicate RateI = 0 and RateII = 0. Q3 (both rates negative) corresponds to two-sided extrusion, points along the negative RateI or RateII axes correspond to strictly one-sided extrusion, and off-diagonal populations indicate one-sided extrusion with slippage.

Stacked bar charts summarize the fractions of two-sided (blue), strictly one-sided (pink), and one-sided extrusion with slippage (green) for each fluorophore subpopulation.

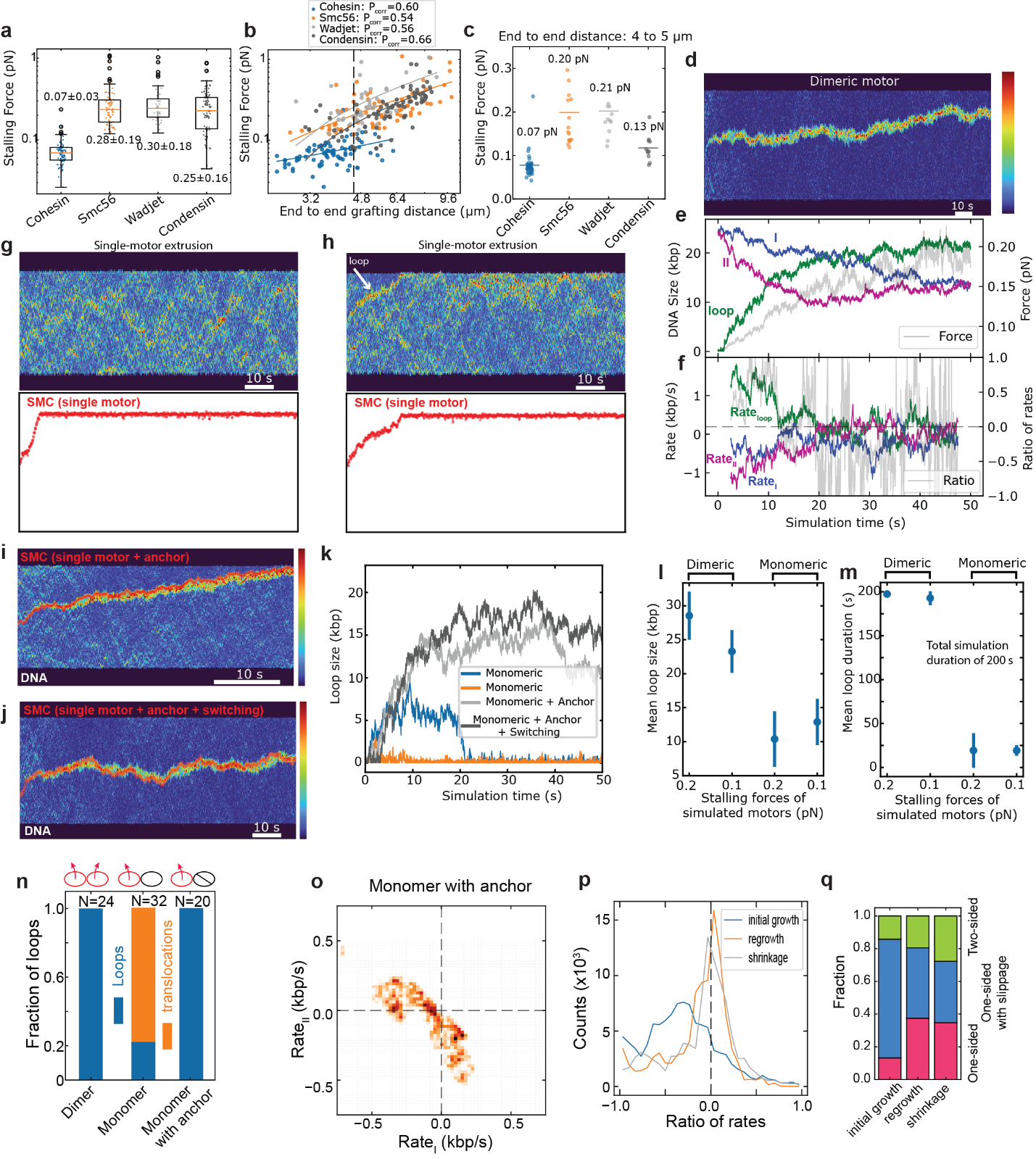

**Supplementary Figure S7. Calibration of simulation parameters and comparison of monomeric and dimeric loop extrusion models.** (a) Distribution of experimentally measured stalling forces for cohesin, Smc5/6, Wadjet, and condensin. Boxes indicate the interquartile range, and whiskers indicate the full distribution.

(b) Relationship between DNA extension and stalling force for the different SMC complexes. Solid lines indicate linear regression fits. The black dashed line marks the 5 µm DNA extension used in simulations.

(c) Stalling forces measured at DNA extensions between 4 and 5 µm. Horizontal bars indicate mean values.

**Dimeric motor simulations.**
(d) Representative kymograph generated from simulations of a dimeric handcuff motor.
(e) Corresponding time traces of DNA segment sizes (loop, I, and II) and DNA tension.
(f) Corresponding RateI, RateII, and ratio-of-rates traces.

**Monomeric motor simulations.**
(g,h) Representative kymographs of monomeric handcuff simulations without stable loop formation (g) and with transient loop formation (h). The arrow in (h) indicates a small loop.

**Monomeric motor simulations with anchoring.**
(i,j) Representative kymographs of monomeric handcuff simulations with an anchoring interaction applied to the non-extruding side, showing examples without (i) and with (j) direction switching.
(k) Representative loop-size trajectories for monomeric simulations without anchoring, with anchoring, and with anchoring plus direction switching.

(l,m) Mean maximum loop sizes (l) and loop lifetimes (m) for dimeric and monomeric simulations without anchoring. Loop lifetimes were measured over a total simulation duration of 200 s.

(n) Fraction of simulation trajectories resulting in stable loop extrusion versus DNA translocation for dimeric (left), monomeric (middle), and anchored monomeric (right) motors.

(o-q) Directionality analysis of anchored monomeric motor simulations, showing the RateI-versus-RateII quadrant plot (o), ratio-of-rates distributions during initial growth, regrowth, and shrinkage phases (p), and the corresponding fractions of two-sided (green), one-sided with slippage (blue), and strictly one-sided (pink) extrusion states (q).

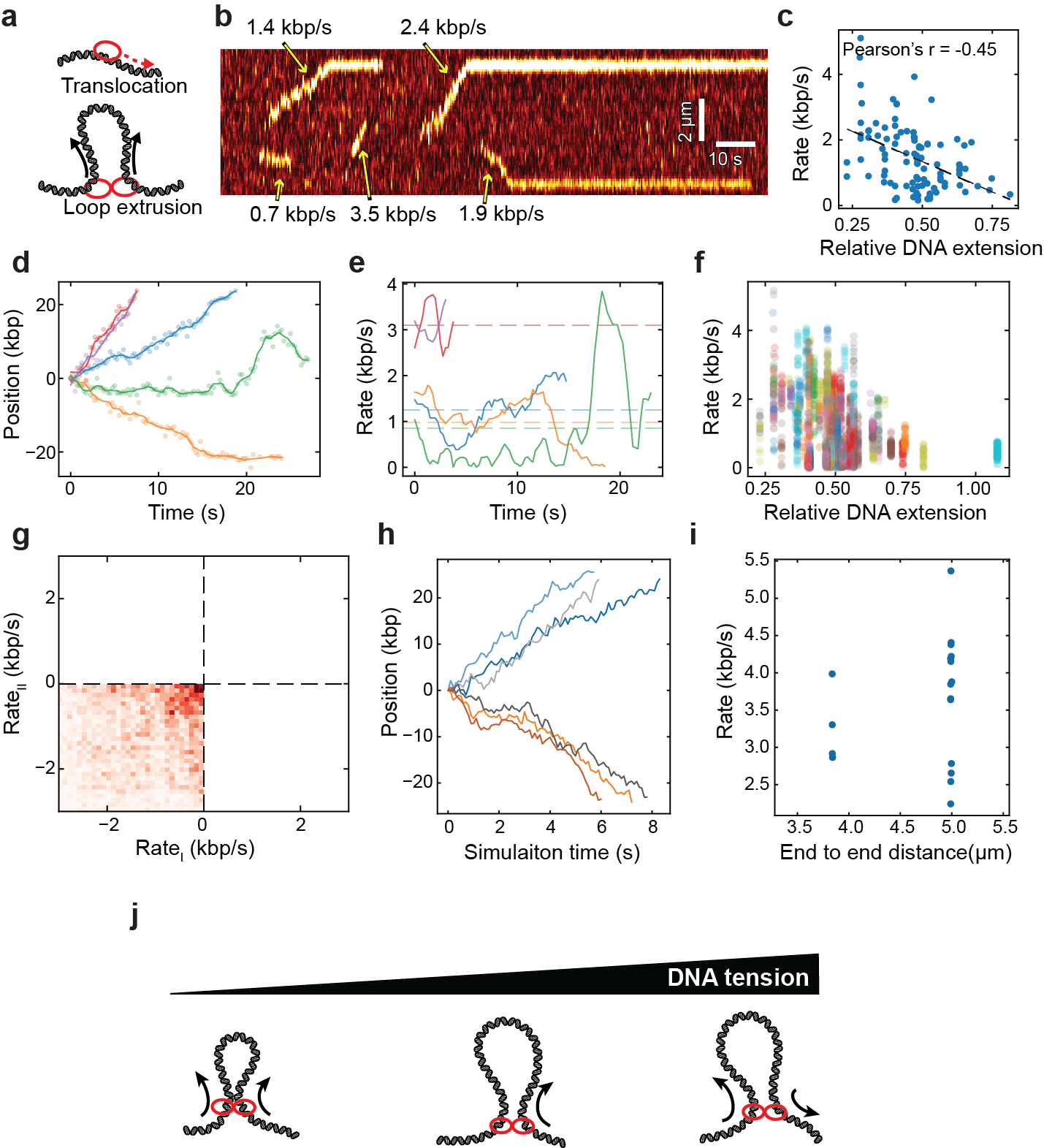

**Supplementary Figure S8. Dimeric SMC complexes comprise two independent translocating motors.**
(a) Schematic illustrating loop extrusion by a dimeric SMC complex (two active motors) and DNA translocation by a monomeric SMC complex (one active motor).

(b) Representative kymograph showing multiple translocation events by Alexa647-labeled octameric Smc5/6 complexes on DNA.

(c) Translocation rate as a function of relative DNA extension, showing a progressive decrease in translocation speed with increasing DNA extension.

(d,e) Representative translocation trajectories (d) and the corresponding instantaneous translocation rates (e), calculated using a 3-s sliding window. Substantial rate variability is observed both between molecules and within individual trajectories.

(f) Scatter plot of translocation rate versus relative DNA extension. Data points from a single Smc5/6 molecule are highlighted, illustrating rate fluctuations within an individual trajectory.

(g) Model RateI-versus-RateII distribution generated by randomly pairing translocation rates measured at the same DNA extension. The resulting distribution recapitulates the broad range of rate combinations observed during loop extrusion.

(h,i) Simulated translocation trajectory of a single-ring handcuff motor (h) and the corresponding distribution of translocation rates (i), demonstrating intrinsic rate variability in the absence of regulatory asymmetry.

.
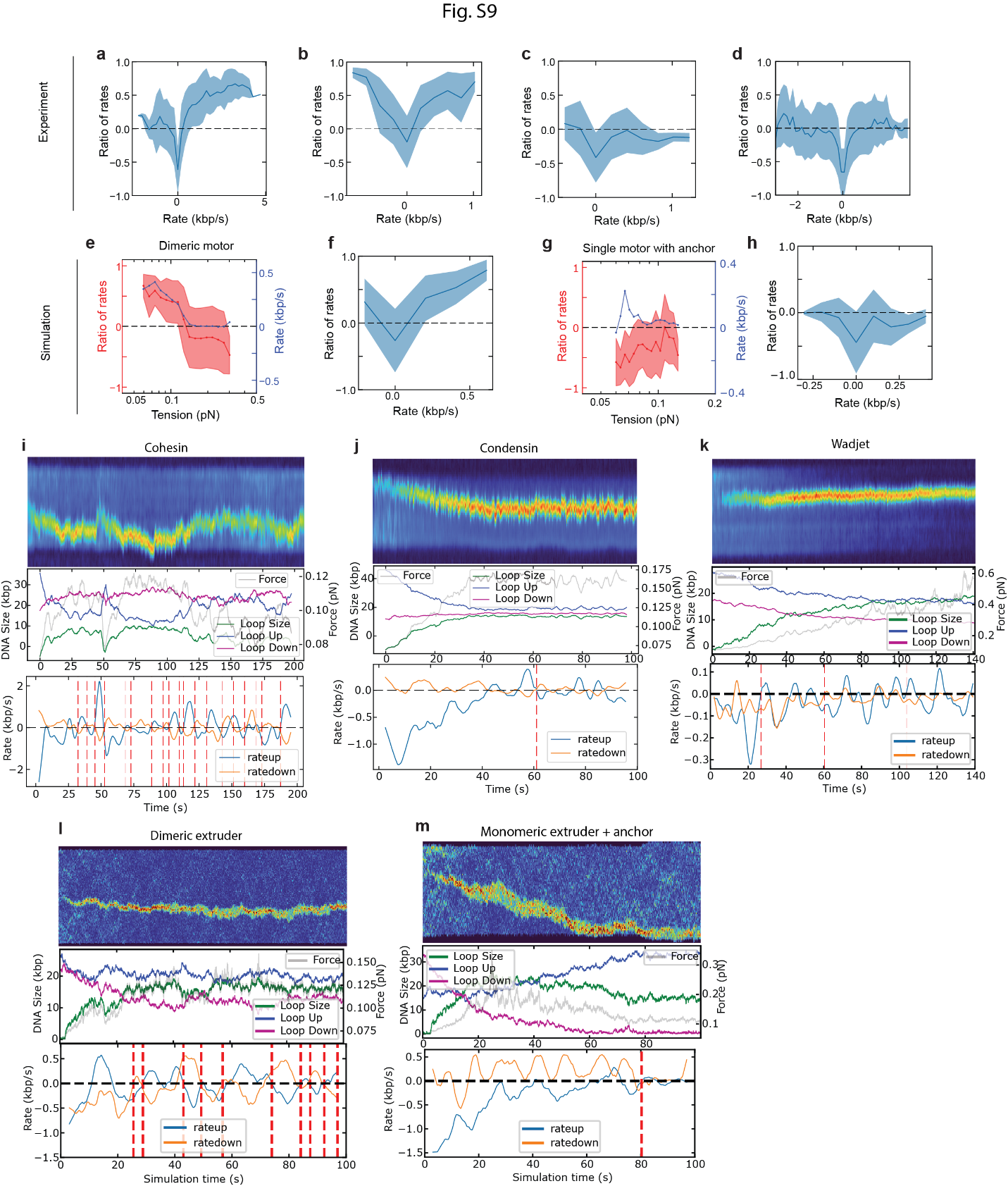

**Supplementary Figure S9. Tension-dependent directionality and direction switching during loop extrusion for experimental and simulated SMC complexes.** (a–d) Ratio-of-rates (mean ± s.d., blue shading) plotted against loop extrusion rate for Smc5/6 (a), Wadjet (b), condensin (c), and cohesin (d).

(e,f) Simulation results for the dimeric motor model. (e) Ratio-of-rates (red, left axis, mean ± s.d.) and loop extrusion rate (blue, right axis) plotted as a function of DNA tension. (f) Ratio-of-rates (mean ± s.d., blue shading) plotted as a function of loop extrusion rate. (g,h) Simulation results for the anchored monomeric motor model. (g) Ratio-of-rates (red, left axis, mean ± s.d.) and loop extrusion rate (blue, right axis) plotted as a function of DNA tension. (h) Ratio-of-rates (mean ± s.d., blue shading) plotted as a function of loop extrusion rate.

(i–k) Experimental data for direction switching during loop extrusion by cohesin (i), condensin (j), and Wadjet (k). Each panel shows, from top to bottom: a representative kymograph, time traces of DNA segment sizes (Loop Size, Loop Up, Loop Down) and DNA tension (grey), and the corresponding Rate_I_ (blue) and Rate_II_ (orange) time traces. Red dashed lines indicate identified direction-switching events.

(l) Simulation results for a dimeric extruder. From top to bottom: simulated kymograph, time traces of DNA segment sizes and tension, and Rate_I_ (blue) and Rate_II_ (orange) traces with red dashed lines indicating spontaneous direction-switching events. (m) Equivalent simulation results for a monomeric extruder with anchoring force.

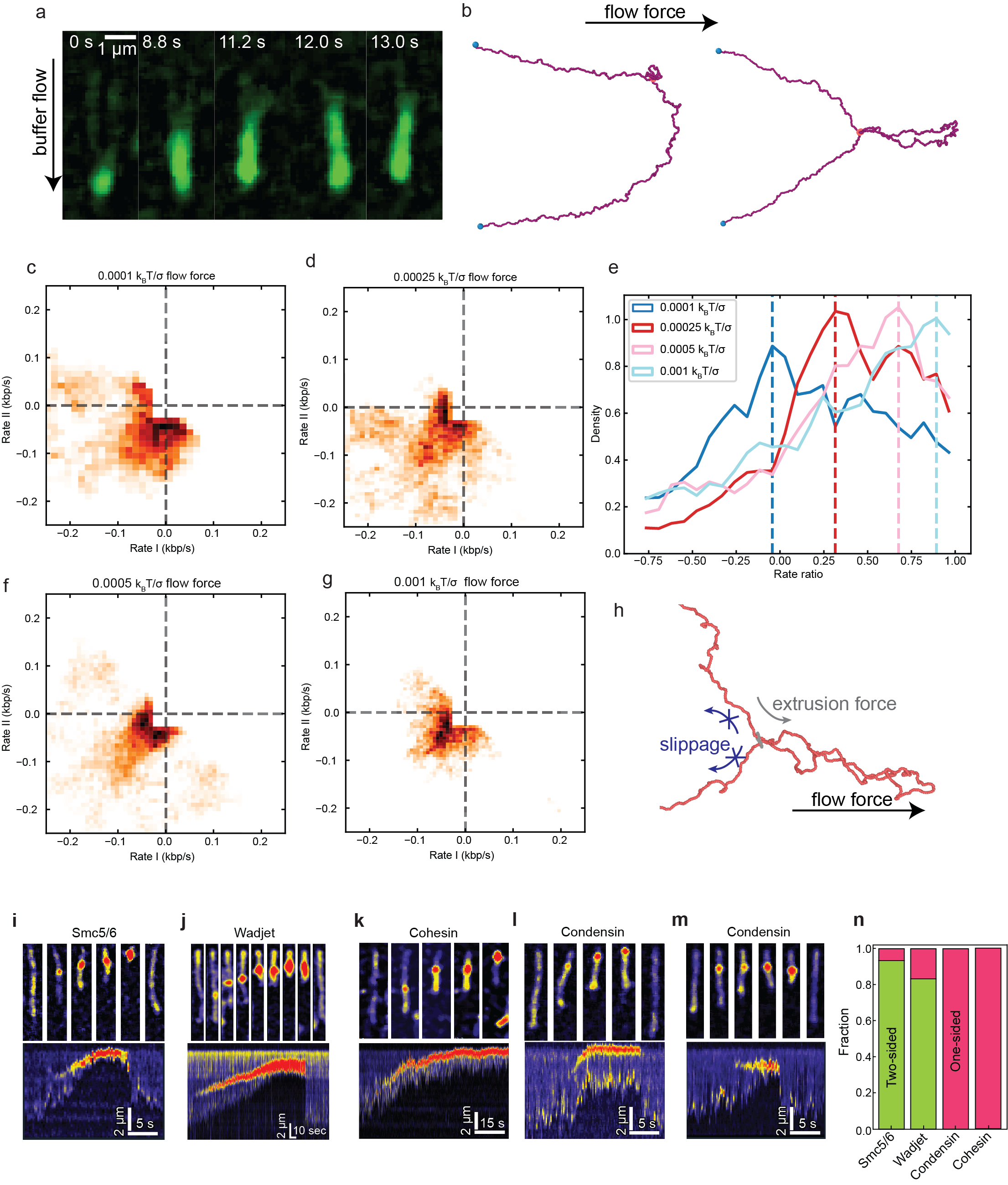

**Supplementary Figure S10. Loop extrusion directionality under side-flow conditions and on single-tethered DNA.** (a) Snapshots of a DNA molecule showing two-sided loop extrusion by Smc5/6 on double-tethered DNA under constant buffer flow. The progressive and symmetric shortening of the DNA from both sides of the loop is indicative of two-sided extrusion. (b) Snapshots of simulated DNA at the start of a simulation (left) and after forming a mature loop (right) under a constant perpendicular flow force. Blue dots indicate the tethered ends of the DNA.

(c–f) Quadrant plots of Rate_I_ versus Rate_II_ during loop growth in simulations with increasing flow forces of 0.0001 k_B_T/σ (b), 0.00025 k_B_T/σ (c), 0.0005 k_B_T/σ (d), and 0.001 k_B_T/σ (e), with extrusion force held constant. As flow force increases, the distribution progressively shifts toward Q3, reflecting a transition to more two-sided extrusion.

(g) Ratio-of-rates density distributions for all four flow forces shown in (b–e). Dashed vertical lines mark the peak of each distribution. Increasing flow force shifts the peak toward positive values, indicating that side-flow promotes two-sided extrusion by counteracting DNA slippage from the passive side of the loop.

(h) Zoomed-in view of a simulated loop under constant flow force, illustrating the competing extrusion force and flow force, with arrows indicating the direction of slippage from the non-extruding side.

(i–m) Loop extrusion on single-tethered DNA, in which one end is anchored to the surface and the other remains free, minimizing tension accumulation during loop growth. Each panel shows snapshots of the DNA molecule (top) and the corresponding kymograph (bottom) for Smc5/6 (i), Wadjet (j), cohesin (k), and condensin at low concentration (l,m).

(n) Fractions of two-sided (green) and one-sided (pink) extrusion quantified for each SMC complex on single-tethered DNA.

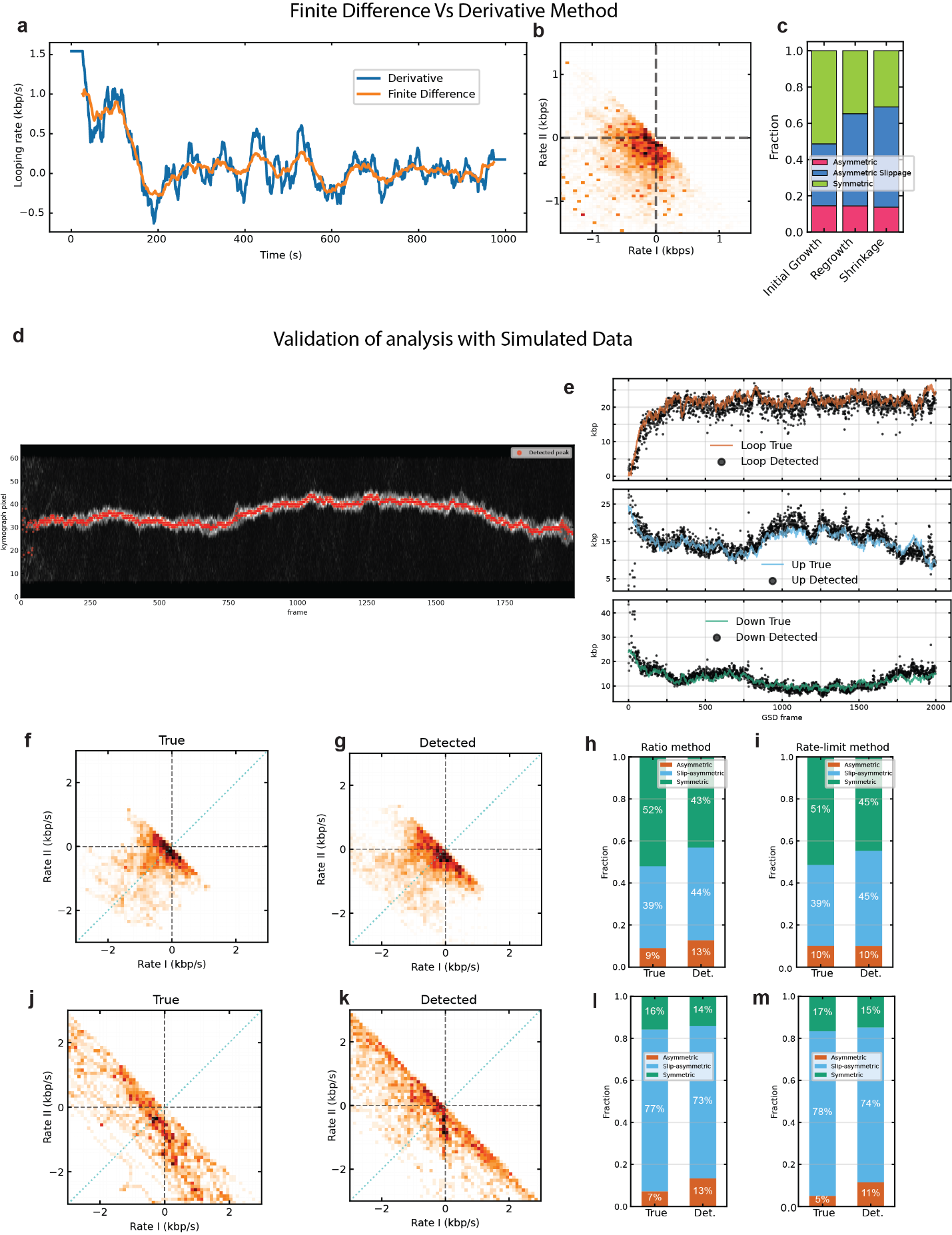

**Supplementary Figure S11. Validation of analysis methods.** (a–c) Comparison of finite difference and derivative methods for calculating loop extrusion rates. (a) Time trace of looping rate for a representative Smc5/6 loop extrusion event, computed using the derivative of the smoothed size trace (blue) and the sliding-window finite difference method (ΔSize/Δt; orange). Both methods yield closely overlapping rate profiles. (b) Quadrant plot of Rate I versus Rate II for Smc5/6 loop extrusion events analyzed using the finite difference method, showing the same distribution pattern as obtained with the derivative approach. (c) Stacked bar chart comparing the fractions of two-sided (green), one-sided with slippage (blue), and strictly one-sided (pink) extrusion across loop extrusion phases (initial growth, regrowth, shrinkage), confirming that the two methods yield equivalent directionality classifications. (d–m) Validation of the loop detection and directionality analysis pipeline using molecular dynamics simulated data with known ground truth. (d) A simulated (MD) kymograph with detected loop peaks overlaid (orange), demonstrating accurate loop identification. (e) Comparison of true (solid lines) and detected (black dots) size traces for the loop region (top), upper flanking segment (middle), and lower flanking segment (bottom), showing close agreement between ground truth and detected values. (f,g) Quadrant plots of Rate_I_ versus Rate_II_ from the true simulated rates (f) and the detected rates from the analysis pipeline (g) for a simulated two-sided extruder. (h,i) Corresponding directionality fractions for the true and detected data, quantified using the ratio method (h) and the rate-limit method (i), showing that both methods faithfully recover the input directionality distribution with only minor deviations. (j,k) Quadrant plots of true (j) and detected (k) rates for a simulated one-sided extruder. (l,m) Corresponding directionality fractions calculated using the ratio method (l) and rate-limit method (m), confirming that the analysis pipeline accurately recovers the predominantly two-sided input distribution. Together, these validations demonstrate that our finite difference rate calculation and loop detection pipeline faithfully capture the directionality of loop extrusion with minimal systematic bias.
